# Age-related decline of the unfolded protein response in the heart promotes protein misfolding and cardiac pathology

**DOI:** 10.1101/2021.06.16.448596

**Authors:** Christoph Hofmann, Erik A. Blackwood, Tobias Jakobi, Clara Sandmann, Julia Groß, Nicole Herzog, Randal J. Kaufman, Hugo A. Katus, Mirko Völkers, Christopher C. Glembotski, Shirin Doroudgar

## Abstract

Cardiac myocyte death during heart failure is particularly detrimental, given that cardiac muscle exhibits limited regenerative potential. Protein aggregation was previously observed in end-stage heart failure, suggesting protein-misfolding in cardiac myocytes as a contributor to the disease process. However, the relationship between protein-misfolding, cardiac myocyte death, and myocardial dysfunction is yet to be clearly established. Here, we showed that protein synthesis and the unfolded protein response (UPR) declined as a function of mammalian postnatal development, especially in tissues with low mitotic activity, such as the heart. A deeper examination in animals models showed that compared to neonatal cardiac myocytes, adult cardiac myocytes expressed lower levels of the adaptive UPR transcription factor, ATF6, as well as lower levels of numerous ATF6-regulated genes, which was associated with susceptibility to ER stress-induced cell death. Further reduction of the ATF6-dependent gene program in ATF6 knock-out mice led to the accumulation of misfolded proteins in the myocardium and impaired myocardial function in response to cardiac stress, indicating that ATF6 plays a critical adaptive role in the setting of cardiac disease. Thus, strategies to increase ATF6 aimed at balancing proteostasis in cardiac myocytes might be a fruitful avenue for the development of novel therapies for heart disease and other age-associated diseases.

**Highlights:**

- The unfolded protein response (UPR) declines as a function of age in adult mammalian tissues with low mitotic activity, such as the heart
- Decreases in the UPR in adult cardiac myocytes is associated with impaired survival during ER stress
- ATF6 loss of function in adult hearts increases protein misfolding and cardiac disfunction during stress

## Introduction

Regulation of proteostasis is critical for maintaining a functional proteome and the prevention of toxic protein misfolding (Klaips et al., 2018). As organisms age, mechanisms for maintaining proteostasis are compromised, which limits the cellular capacity to sufficiently prevent protein misfolding during stress (Kaushik and Cuervo, 2015). This phenomenon has been studied for neurodegenerative diseases in which an age-associated decline in protein quality control and subsequent protein misfolding are major risk factors for disease development and progression (Hipp et al., 2019). The unfolded protein response (UPR) contributes to maintaining protein homeostasis (proteostasis) and was previously shown to decline with age in invertebrates (Ben-Zvi et al., 2009; Schindelin et al., 2012; Taylor and Dillin, 2013). However, whether the UPR declines with age in mammals, and if so, whether this decline contributes to pathology remains unresolved.

Proteostasis is especially important in tissues with low mitotic activity, such as the brain and heart, partly due to the lack of asymmetric inheritance of protein aggregates during cell division which can reduce the amount of misfolded proteins in the daughter cell (Aguilaniu et al., 2003; Hipp et al., 2019). Previously, protein aggregation has been observed during cardiac aging and in end-stage heart failure (Ayyadevara et al., 2016; Mohammed et al., 2014; Okada et al., 2004; Rainer et al., 2018; Sanbe et al., 2004; Tannous et al., 2008) and misfolded proteins are known to directly impair cardiac function (Hofmann et al., 2019; Pattison et al., 2008; Sanbe et al., 2004; Wang et al., 2001). However, whether an age-related decline of the UPR contributes to cardiac protein misfolding and impairs myocardial function during times of cardiac stress are not known.

In metazoans, the UPR initiates transcriptional and translational cellular reprogramming in response to ER stress. The three ER transmembrane proteins, inositol requiring enzyme 1α/β (IRE1) (Cox et al., 1993; Morl et al., 1993), PKR-like ER kinase (PERK) (Harding et al., 1999), and activating transcription factor 6α/β (ATF6) (Haze et al., 1999) regulate three branches of the UPR aimed at restoring proteostasis. ATF6 differs significantly from PERK and IRE1 in terms of its mechanism of activation. Upon accumulation of misfolded proteins, ATF6 transits to the Golgi apparatus where it is cleaved by site-specific proteases S1P and S2P (Haze et al., 1999; Shen et al., 2002). Active ATF6 translocates to the nucleus, where it serves as a potent and short-lived transcription factor, inducing adaptive proteins that restore ER folding capacity and resolve ER stress (Glembotski, 2014; Haze et al., 1999). Previously, ATF6 was shown to be essential for cardiac function, especially during pathological conditions that are associated with proteotoxic stress, such as cardiac growth or reperfusion (Blackwood et al., 2019a; Jin et al., 2017; Martindale et al., 2006). Therefore, while it has not yet been studied, an age-related decline in the ATF6 branch of the UPR would be predicted to promote myocardial dysfunction.

## Results

### Impairment of UPR-mediated protein quality control as a function of age

We first assessed the protein synthesis aspect of proteostasis, as well as the expression of several UPR-regulated gene products during postnatal development in different tissues in neonatal (one day old), young adult (10 weeks old), and middle-aged (52 weeks old) mice. To measure changes in protein synthesis rates, *in vivo*, mice were injected with puromycin. Detection of puromycin-labelled peptides by immunoblotting using an anti-puromycin antibody showed age-associated decreases in protein synthesis rates in the tissues of low mitotic activity heart, skeletal muscle, and brain (**Fig. 1A-C**). While we observed a significant age-dependent reduction of protein synthesis in mitotic tissues such as kidney and liver, this reduction was less pronounced compared to that in tissues with low mitotic activity (**Fig. 1D, E**). We also found a reduced abundance of the canonical hallmark adaptive UPR-induced gene products, chaperones 78 kDa glucose-regulated protein (GRP78) and GRP94, and protein disulfide-isomerase A6 (PDIA6), in young adult and middle-aged mice tissues compared to neonatal, a reduction that was more pronounced in low-mitotic tissues (**Fig. 1F-J**). Importantly, we found a highly significant correlation between protein synthesis and UPR gene expression in all samples, which may indicate protein synthesis and the UPR are coordinately regulated (**Fig. 1K**).

**Figure 1.**
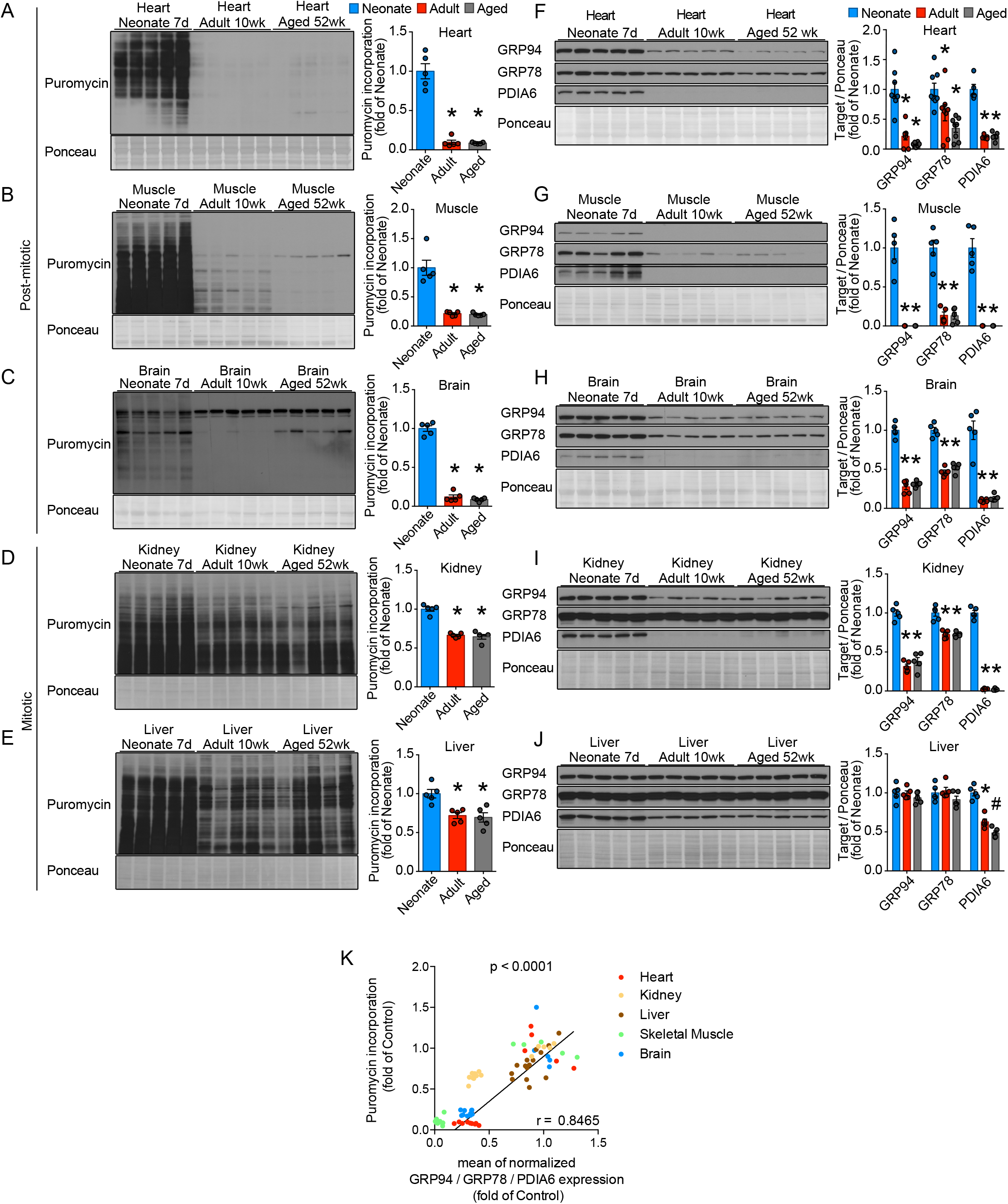
Extensive downregulation of protein synthesis and the UPR in tissues with low mitotic activity during mammalian adulthood. Immunoblots of puromycin and respective quantifications for the low-mitotic tissues heart (A), skeletal muscle (B), brain (C) and the mitotic tissues kidney (D) and liver (E). Immunoblots of the UPR gene products GRP94, GRP78, and PDIA6 for the low-mitotic tissues heart (F), skeletal muscle (G), brain (H) and the mitotic tissues kidney (I) and liver (J). (K) Scatter plot showing the correlation between puromycin incorporation and the mean of the normalized expression of GRP94, GRP78, and PDIA6. Colors for each organ are shown on the right. r was calculated with GraphPad Prism v7.0. * indicates p<0.05 from neonate; # indicates p<0.05 from adult.

### Reduced responsiveness of the UPR in the adult heart makes cardiac myocytes susceptible to proteotoxic stress

To more broadly assess the pathophysiological importance of the UPR aspect of proteostasis in the heart as a function of age, we examined human heart from different ages ranging from newborn to adult (Cardoso-Moreira et al., 2019) for changes in the expression of canonical UPR genes during human postnatal development and aging. We observed an age-related decline in the expression of these UPR genes from young adulthood in human hearts (**Fig. 2A**).

**Figure 2.**
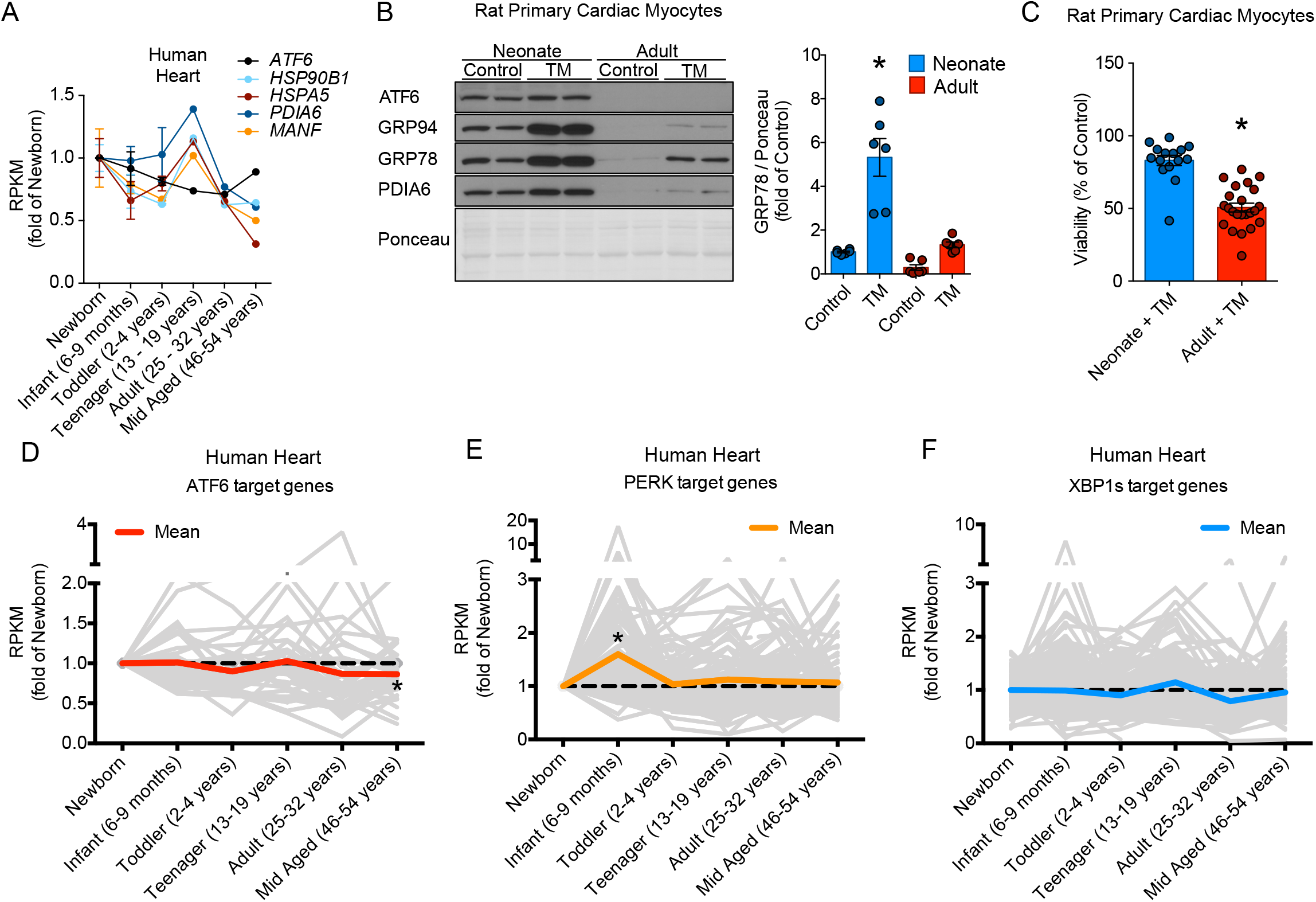
Impaired expression and responsiveness of the UPR in the adult heart. (A) Normalized RPKM values of the respective genes for human RNA sequencing data of the indicated age groups; newborn n=2, infant n=3, toddler n=2, teenager n=1, young adult n=1, older mid-aged n=1. (B) Immunoblots and quantifications for ATF6, GRP94, GRP78, and PDIA6 of neonatal and adult rat primary cardiac myocytes treated with vehicle (DMSO; control) or TM for 24h. (C) Relative viability of neonatal and adult rat cardiac myocytes after 24h of 20μg/mL TM treatment. Values are normalized to the viability of vehicle (DMSO) treated cells of the same experiment. Each dot represents the viability of all cells in a random 10× magnified cell culture view. Data are representative of three independent experiments. * indicates p<0.05 from neonate. (D-F) Normalized RPKM values of the target genes of ATF6, PERK, and XBP1 pathways in human RNA sequencing data of the indicated age groups. * indicates p<0.05 of mean RPKM from newborn.

While myocytes comprise more than 90% of the mass of the heart, they do not represent the dominant cell type, as about 70% of cardiac cells are nonmyocytes, such as endothelial cells and fibroblasts, which maintain the ability to divide throughout development (Pinto et al., 2016). Accordingly, to characterize the UPR in cardiac myocytes as a function of age we isolated ventricular myocytes from neonatal and adult rats. Similar to data from whole left ventricular lysates, protein synthesis and levels of adaptive UPR-regulated gene products decreased as a function of age in isolated cardiac myocytes (**Fig. 2B; Fig. S1A, C**). However, the decline in UPR gene expression was more pronounced in isolated cardiac myocytes compared to heart extracts, falling to nearly undetectable levels in cardiac myocytes of adult rats (**Fig. 2B; Fig. S1A-C**). While the protein glycosylation inhibitor, tunicamycin (TM) was able to induce UPR genes in adult cardiac myocytes (**Fig. S2**), UPR gene expression did not reach the levels of neonatal cardiac myocytes, even after TM treatment (**Fig. 2B**). Importantly, the decline of the UPR in adult cardiac myocytes was accompanied by impaired survival of adult cardiac myocytes in response to TM treatment (**Fig. 2C**), indicating that the reduced UPR responsiveness in adult cardiac myocytes is associated with impaired survival during ER stress. To assess the UPR on a global level, we examined the expression of genes downstream of the all three UPR branches, ATF6, PERK, and IRE1 during human postnatal development and aging (Cardoso-Moreira et al., 2019) and found a significant downregulation of ATF6 genes in mid-age in human heart (**Fig. 2D-F**).

### Decline of ATF6 and the ATF6-regulated UPR gene program with age has negligible effects in the heart at baseline

Since we found that ATF6-dependent target genes decrease in mid-age in the human heart, we focused on the ATF6 branch of the UPR. We hypothesized that the reduced expression of ATF6 in young adult and middle-aged mice is at least partly responsible for the decreased expression of UPR target genes in the heart. ATF6 knockout (KO) mice develop normally throughout adulthood and exhibit no overt cardiac phenotype (Wu et al., 2007; Yamamoto et al., 2007), indicating that ATF6 is not essential for normal cardiac development and function under non-stressed conditions (Jin et al., 2017). We performed transcript profiling of hearts from ATF6 KO mice. The absence of ATF6 was confirmed by RT-PCR (Fig. 3A). While RNA-sequencing revealed that “Response to endoplasmic reticulum stress” was among the most regulated signaling pathways in adult ATF6 KO mouse hearts (Fig. 3B), only a few UPR genes were significantly affected in ATF6 KO hearts compared to WT hearts (Fig. 3C, D), which is in agreement with previous studies that showed no major consequence of ATF6 deficiency on cardiac myocyte survival or UPR gene expression under baseline conditions (Correll et al., 2019; Jin et al., 2017; Wu et al., 2007; Yamamoto et al., 2007). ATF6 controls the expression of target genes by binding cis-acting promoter elements, such as the ER stress response element (ERSE) (Yamamoto et al., 2004). However, activation of the IRE1 branch of the UPR results in the cytoplasmic RNA splicing of uXBP1 to spliced XBP1, which when translated generates a transcription factor that binds to the same and additional promoter elements (Yamamoto et al., 2004). Therefore, expression of ATF6 target genes during ATF6 deficiency might be affected by activation of XBP1. Indeed, pharmacological inhibition of ATF6 activation with PF-429242 rapidly induced XPB1 splicing (**Fig. S3**), indicating that ATF6 deficiency in cardiac myocytes may be at least partly compensated by IRE1 pathway activation. Taken together, reduced ATF6 expression during cardiac aging seems to be at least partly responsible for the reduced UPR capacity of adult cardiac myocytes.

**Figure 3.**
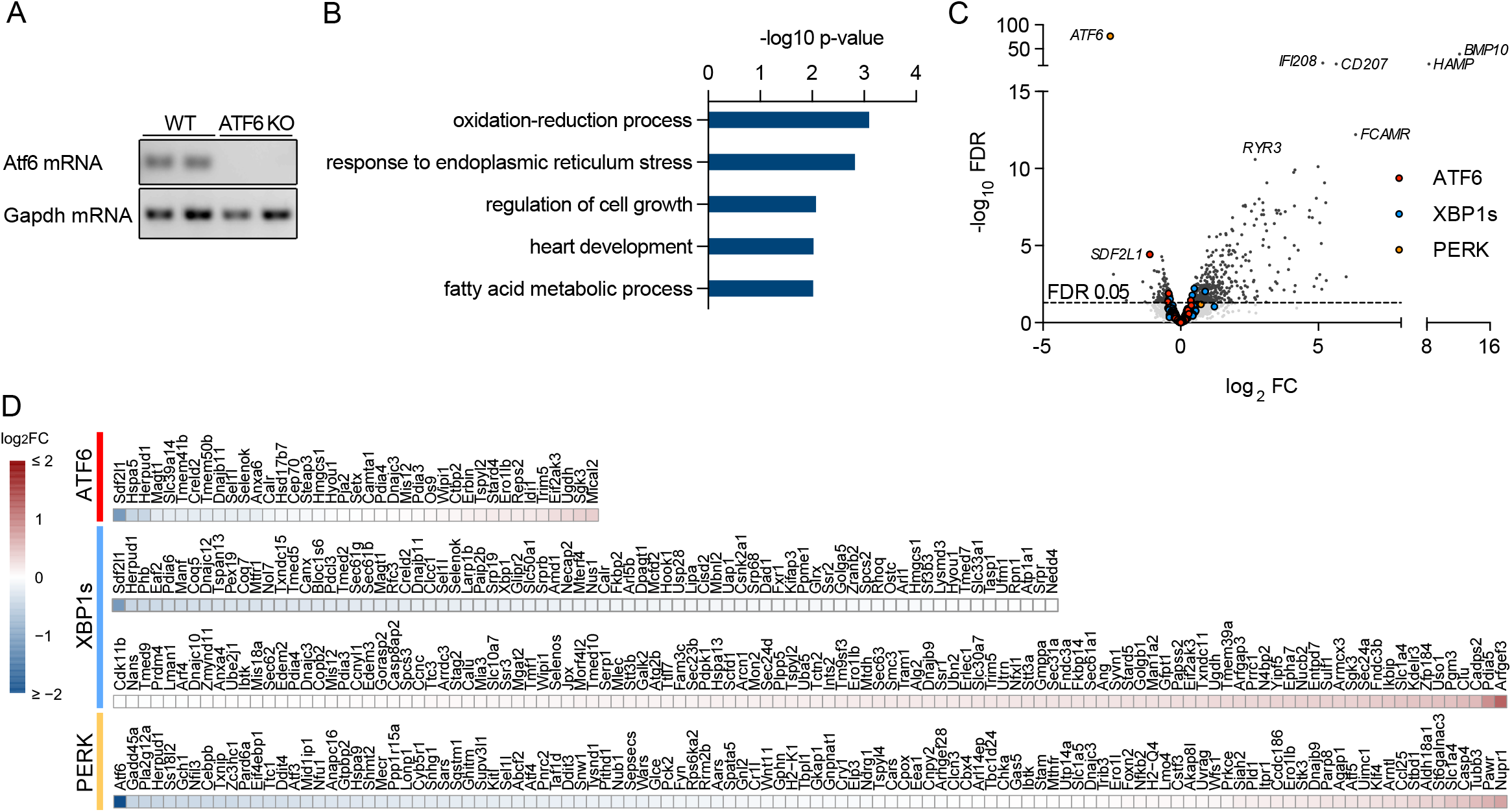
ATF6 protects from ER stress-induced cell death in cardiac myocytes but has only minor impact on cardiac UPR in the absence of stress. (A) ATF6 PCR of WT and ATF6 KO mice showing the successful knock-out of ATF6. (B) Enrichment of gene ontology terms (biological process) of significantly changed genes. (C) Volcano plot of transcriptional changes measured by RNA sequencing in adult ATF6 KO mice compared to WT mice. The highlighted colors indicate ATF6 (red), XBP1s (blue), or PERK (yellow) target genes. The horizontal dotted line indicates an FDR of 0.05. Genes above the line are considered significant (FDR < 0.05). 4 WT and 3 ATF6 KO mice were used for the analysis. (D) Heat map showing the regulation of all ATF6 (red), XBP1s (blue) or PERK (yellow) target genes in ATF6 KO mice.

### Importance of the ATF6 branch of the UPR in adult cardiac myocyte hypertrophy

Various cardiac pathologies are associated with increased protein synthesis, most notably cardiac hypertrophy, in which the increased synthesis of new protein drives the hypertrophic growth of cardiac myocytes (Frey and Olson, 2003). Cardiac myocyte hypertrophy during pathology, while initially adaptive, can eventually lead to impaired ventricular relaxation, filling, and eventually cardiac failure (Frey and Olson, 2003; Hill and Olson, 2008; Tardiff, 2006). Previously it was shown that misfolded proteins accumulate during pathological cardiac hypertrophy, however it is not known which factors contribute to this accumulation, nor is it known whether misfolded proteins contribute to the morbidity of cardiac hypertrophy (Singh and Robbins, 2018). Therefore, we investigated the role of UPR and ATF6 in proteostasis in the setting of pathological cardiac myocyte hypertrophy.

We first analyzed the UPR on an integrated transcriptome, translatome, and proteome dataset of the adult myocardial gene response during cardiac hypertrophy (Doroudgar et al., 2019). Interestingly, a biological process component category analysis revealed that “protein folding” and “response to ER stress” were among the most regulated gene ontology terms in the early phase of cardiac hypertrophy in mouse hearts 2 days after transverse aortic constriction (TAC) (**Fig. 4A-C**), a common surgical model of pressure overload-induced cardiac hypertrophy (Rockman et al., 1991). Several genes that belong to the gene ontology groups “response to unfolded protein”, “endoplasmic reticulum unfolded protein response”, and “response to endoplasmic reticulum stress”, which we merged into the term “unfolded protein response genes”, were found to be strongly regulated both on the transcriptional and translational levels (**Fig. 4D, E**). A more in-depth analysis revealed that this group included genes that are regulated by all three branches of the UPR (**Fig. S4, S5**). The cardiac UPR was only mildly upregulated at 3 hours, peaked at 2 days, and declined again, especially at the transcriptional level at 2 weeks after TAC (**Fig. S4, S5**). To expand our profiling results, we analyzed the ATF6-mediated UPR in isolated adult cardiac myocytes during adrenergic stimulation-induced hypertrophy (**Fig. 4F**). Adult cardiac myocytes exhibited an increase of ATF6 levels after adrenergic stimulation (**Fig. 4G**) leading to an increased expression of ATF6 target genes in a dose-dependent manner (**Fig. 4H-J**). Similarly, at 2 days after TAC surgery, when protein synthesis was increased (**Fig. 4K, L**), protein levels of the ATF6 target genes GRP94, GRP78, and PDIA6 were found significantly upregulated (**Fig. 4M**). Taken together, these results indicate that the UPR is upregulated during early phases of pathological hypertrophic growth in cardiac myocytes of the adult heart.

**Figure 4.**
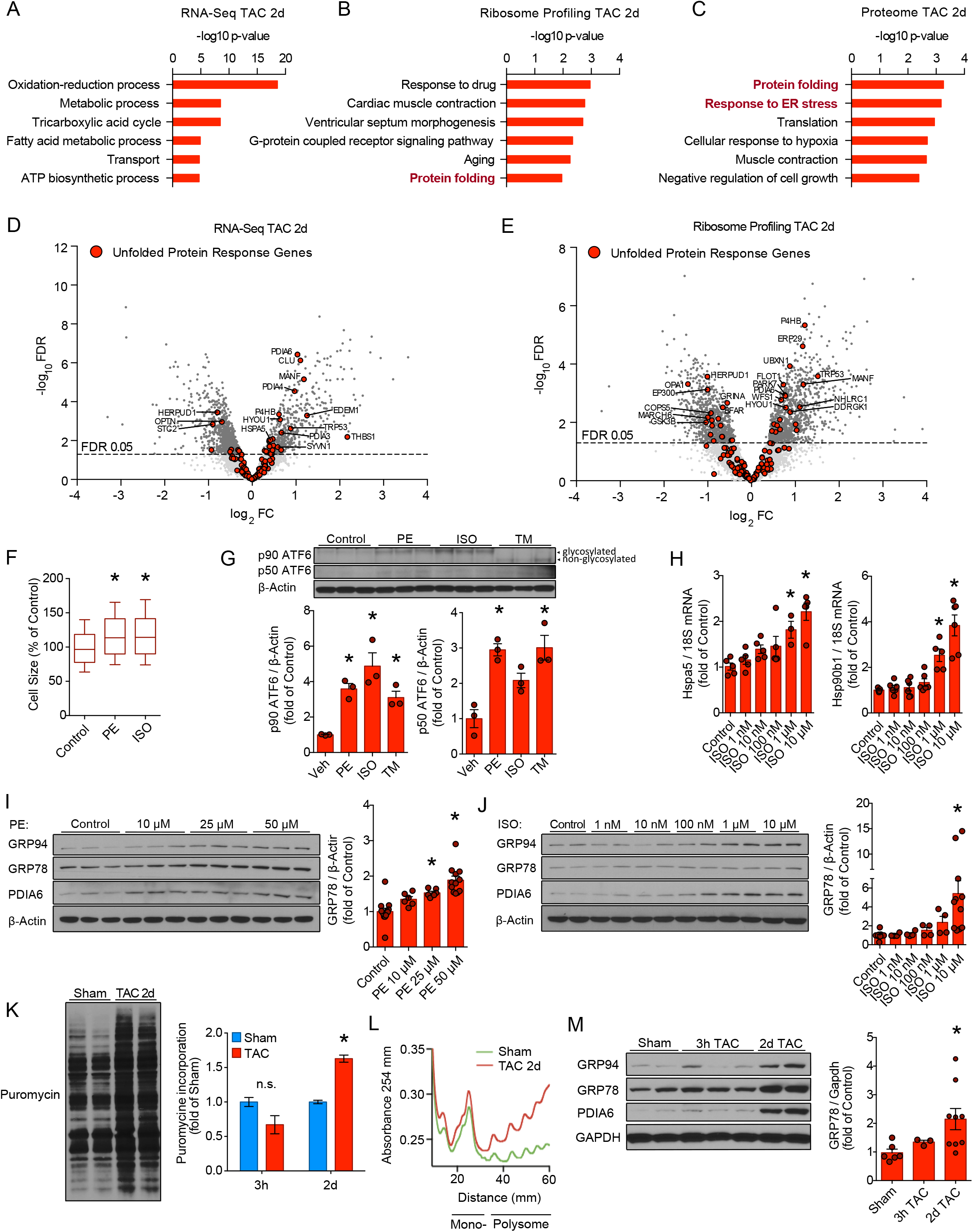
Induction of the UPR during early adult cardiac hypertrophy. (A-C) Enrichment of gene ontology terms (biological process) of RNA-Seq, ribosome profiling, and quantitative mass spectrometry-based proteomics 2d post-TAC or sham surgery. RNA-Seq and ribosome profiling libraries were derived from left ventricular lysates of Ribo-Tag mice, proteome data is based on isolated cardiac myocytes from similarly treated mice. (D and E) Transcriptional or translational regulation of UPR genes 2d after TAC surgery. Transcripts were considered significant when false discovery rate (FDR) was <0.05. (F) Cell surface area measurements of adult cardiac myocytes in response to PBS (Veh), phenylephrine (PE), or isoproterenol (ISO) for 24h. At least 400 cells were counted in total per condition in three independent experiments. (G) p90 ATF6 immunoblots and respective quantification of adult cardiac myocytes after 24h vehicle, PE, ISO, or TM treatment. The two visible bands of p90 ATF6 represent its glycosylated and non-glycosylated form after treatment with the glycosylation inhibitor TM. (H) mRNA levels of GRP78 (Hspa5) or GRP94 (Hsp90b1) after 24h treatment of adult cardiac myocytes with increasing PE or ISO dosages normalized to 18S mRNA. (I and J) Immunoblots and respective quantifications of GRP94, GRP78, and PDIA6 in adult cardiac myocytes treated with increasing dosages of Veh, PE, or ISO for 24h, normalized to β-actin protein levels. (K) Representative puromycin immunoblot of left ventricular lysates of sham and 2d TAC treated mice with quantifications of mice after 3h and 2d TAC normalized to their respective sham control. (L) Polysome profiles of 10 weeks old mouse left ventricular lysates 2d after sham or TAC surgery. (M) Immunoblots and respective quantification of GRP94, GRP78, and PDIA6 of left ventricular lysates from mice 3h or 2d after TAC surgery. Sham mice were sacrificed after 2d. * indicates p<0.05 from control.

### Induction of the ATF6-mediated UPR is necessary to prevent protein misfolding during adult cardiac hypertrophy

After establishing that ATF6 is activated in cardiac myocytes during cellular growth, we examined the involvement of ATF6 in cardiac hypertrophy. Inhibition of ATF6 in neonatal cardiac myocytes by siRNA-mediated knock-down attenuated neonatal cardiac myocyte hypertrophy in response to PE (**Fig. 5A, B**). Next, adult WT and ATF6 KO mice were subjected to implantation of isoproterenol (ISO)-loaded osmotic mini pumps or TAC surgery, two *in vivo* models of cardiac hypertrophy, and examined after 7 days, at a time point when cardiac protein synthesis and hypertrophic growth reach maximum (Wang et al., 2017). ISO and TAC both increased heart weight of WT mice, an effect that was impaired in ATF6 KO mice (**Fig. 5C, E**). Additionally, cardiac function, assessed by transthoracic echocardiography, was significantly impaired at 7 days in ATF6 KO mice, a time point when cardiac function remains preserved in WT mice (**Supp. Table I, II; Fig. 5D, F**) (Takaoka et al., 2002). ATF6 KO hearts from ISO-treated mice showed increased expression of pathologic cardiac remodeling marker genes *Nppa*, *Nppb*, *Myh7,* and *Col1a1* and lower transcript levels of the Ca^2+^ handling gene product *Atp2a2*, compared to WT hearts (**Fig. 5G, H**). These markers of pathologic cardiac remodeling were not significantly different in ATF6 KO mice compared to WT mice after TAC surgery (**Fig. 5G, H**). To examine whether changes in the functional outcome of ATF6 KO mice are due to the loss of ATF6 transcriptional activity and its adaptive gene program, we expressed an adenovirus encoding a dominant negative form of ATF6, dnATF6 (Thuerauf et al., 2001). Furthermore, PF-429242, a chemical inhibitor of ATF6 activation was examined. Both dnATF6 and PF-429242 significantly reduced cardiac myocyte hypertrophy (**Fig. 5I-K**).

**Figure 5.**
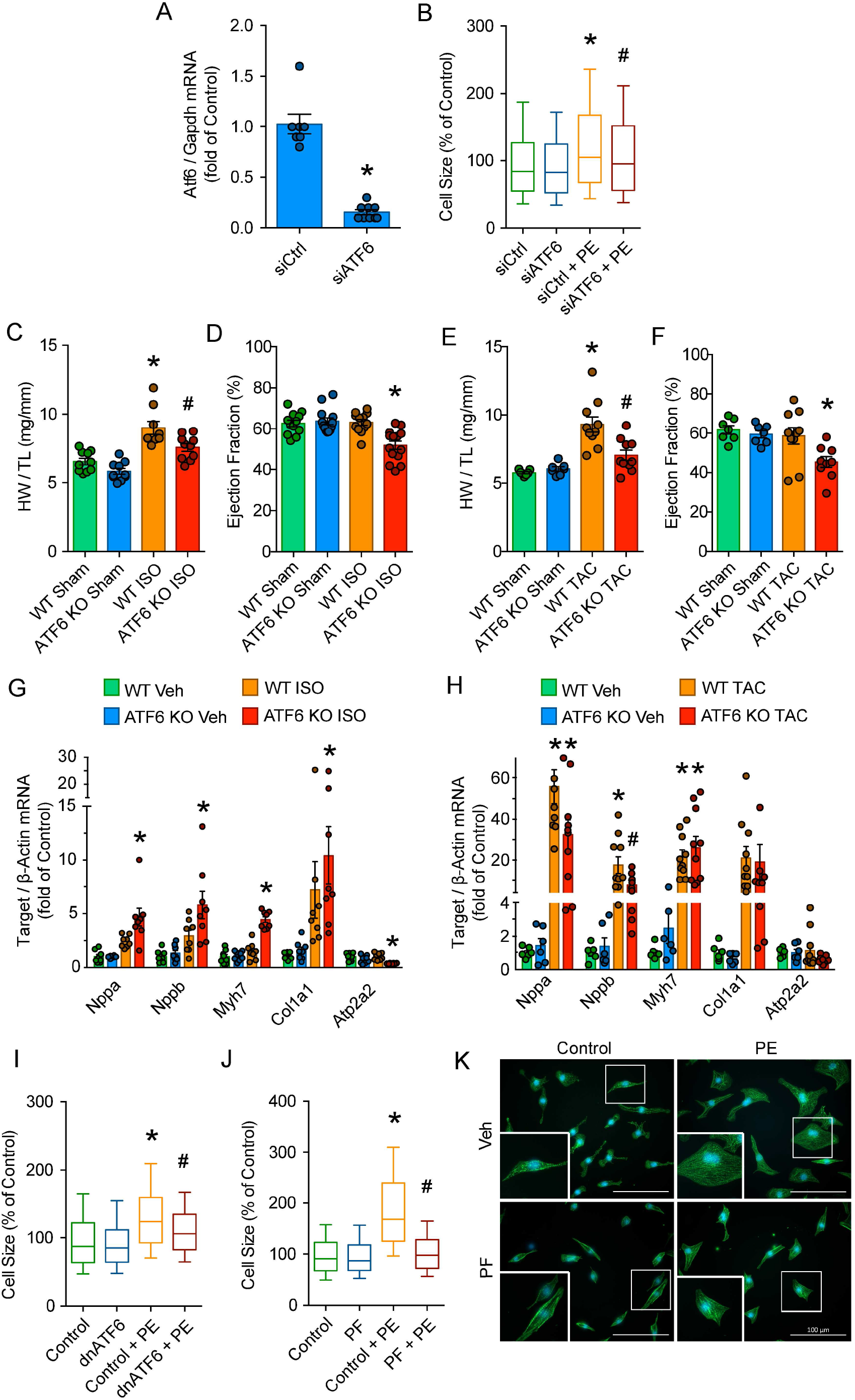
ATF6 insufficiency impairs cardiac hypertrophy in response to stress. (A) ATF6 mRNA levels after siRNA mediated knock-down in neonatal cardiac myocytes. (B) Cell surface area measurements of neonatal cardiac myocytes after siRNA-mediated ATF6 knock-down in response to PE for 24h. (C and E) Heart weight (HW) to body weight (BW) ratio of WT and ATF6 KO mice subjected to sham or ISO pump implantation, or sham or TAC surgeries. (D and F) Ejection fraction of WT and ATF6 KO mice subjected to sham or ISO pump implantation, or sham or TAC surgeries. (G) and (H) mRNA levels of respective maladaptive remodeling genes normalized to β-Actin of mice subjected to sham or ISO pump implantation, or sham or TAC surgeries. (I) Cell surface area measurements of neonatal cardiac myocytes after dnATF6 treatment in response to PE for 24h. (J) Cell surface area measurements and representative immunofluorescence images (K) of neonatal cardiac myocytes after PF-429242 treatment in response to PE for 24h. At least 400 cells were counted in total per condition in three independent experiments for (B), (I) and (J). Samples for (C - H) were collected seven days after treatment. * indicates p<0.05 from control; # indicates p<0.05 from control + PE (B, I and J), from WT Iso (C) or from WT TAC (D and H).

Finally, we examined the mechanisms of the blunted cardiac hypertrophic growth response and subsequent myocardial dysfunction in ATF6 KO mice. Cardiac hypertrophy was coupled with ATF6 activation after TAC surgery, as evidenced by the formation of the active 50kD ATF6, which was completely absent in ATF6 KO mice subjected to TAC surgery (**Fig. 6A**). Importantly, ATF6 KO hearts did not upregulate key UPR protein quality control regulators in response to ISO or TAC, indicating that ATF6 is necessary for UPR induction during cardiac stress (**Fig. 6B, C**). Together with the impaired activation of the UPR, ATF6 mice accumulated poly-ubiquitylated proteins 7 days after TAC surgery, a phenomenon that was only marginally observed in WT mice (**Fig. 6D**). Therefore, we investigated whether cardiac protein misfolding can be directly detected 7 days after TAC surgery when protein synthesis and hypertrophic growth reach maximum. We used immunoblotting with A11, an antibody that detects a common epitope of soluble amyloid oligomers, the entity that is thought to at least partly constitute the pathogenic species of misfolded proteins in cardiac disease (Wang and Robbins, 2006). We observed the accumulation of A11-positive staining in the early phase of TAC-induced cardiac hypertrophy but not in control, sham WT mice (**Fig. 6E**). While ATF6 KO mice did not exhibit detectible misfolded protein aggregates under baseline conditions, amyloid oligomers were increased in ATF6 KO mice after TAC surgery (**Fig. 6E**), demonstrating that ATF6 is essential for preventing the accumulation of misfolded proteins during the early phase of cardiac hypertrophy when protein synthesis is increased but myocardial function is still preserved. This suggests that the accumulation of misfolded proteins occurs during cardiac disease progression, supporting the notion that misfolded proteins are a cause of cardiac dysfunction, not a consequence.

**Figure 6.**
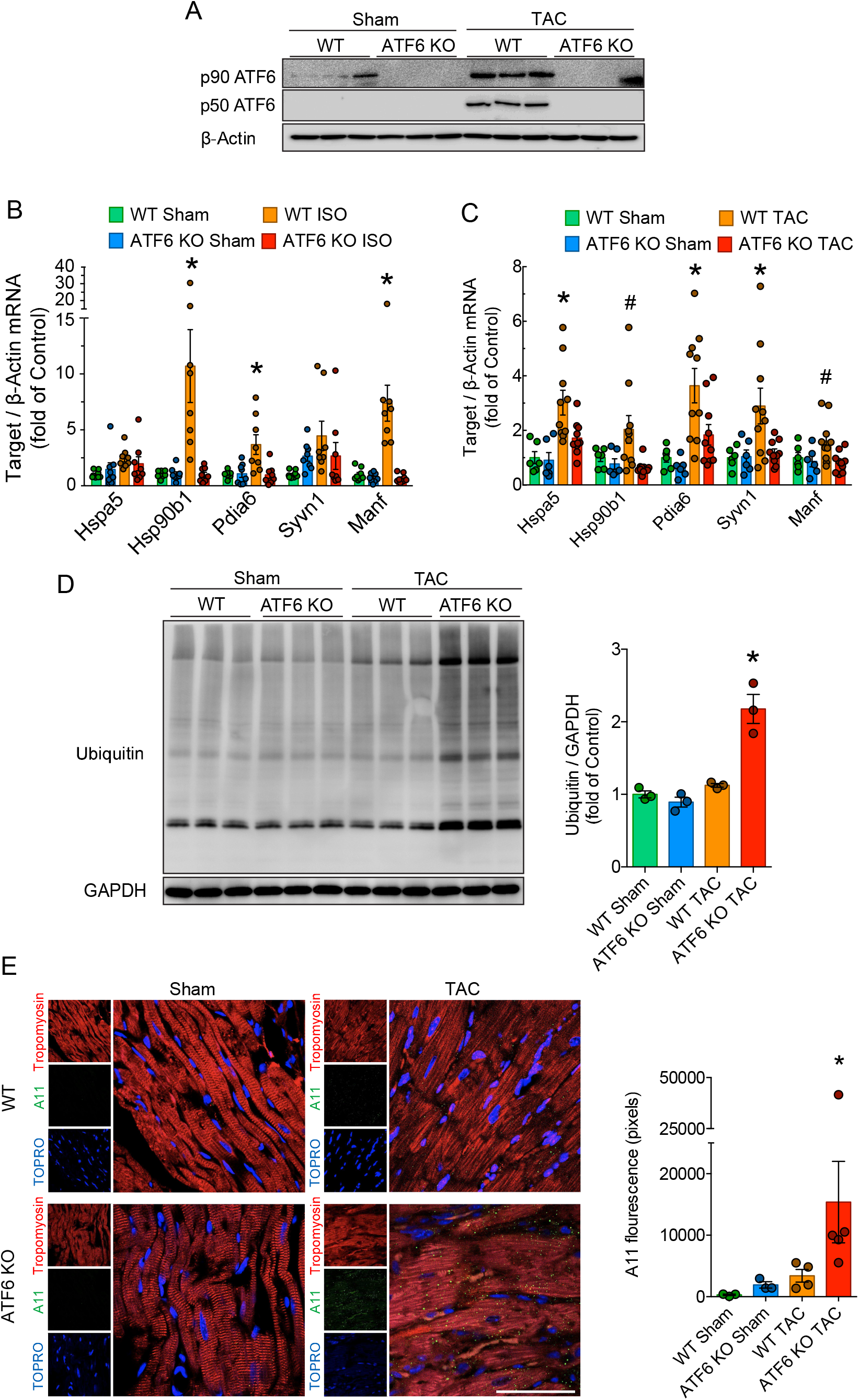
ATF6 deficiency blunts UPR induction in the acute phase of cardiac hypertrophy and leads to myocardial accumulation of misfolded proteins. (A) p90 and p50 ATF6 immunoblots of WT and ATF6 KO mice subjected to sham and TAC surgeries. (B and C) mRNA levels of respective UPR genes normalized to β-Actin of WT and ATF6 KO mice subjected to sham or ISO pump implantation, or sham or TAC surgeries. (D) Ubiquitylated protein immunoblot and respective quantification of WT and ATF6 KO mice subjected to sham and TAC surgeries. (E) A11 immunoflourescence images and respective quantification of left ventricular sections from WT and ATF6 KO mice subjected to sham or TAC surgeries. Sections were co-stained with tropomyosin marking cardiac myocytes and TOPRO-3. Samples for (A - E) were collected seven days after treatment. * indicates p<0.05 from all other conditions; # indicates p<0.05 from control.

## Discussion

In invertebrates the decline in protein homeostasis declines as a function of age in invertebrates contributes to disease-like phenotypes (Kikis et al., 2010). However, analyses of the proteostasis network in aging mammals are limited (Kaushik and Cuervo, 2015). Here, we investigated protein synthesis and UPR-mediated protein quality control capacity during postnatal development and aging in mice and rats. We observed coordinate decreases of the ATF6 branch of the UPR and protein synthesis as a function of age, especially in postmitotic organs, such as the heart. A focused examination of the heart revealed that the decline of the UPR renders adult cardiac myocytes more susceptible to proteotoxic stress, which was at least partly attributable to decreased activity of the ATF6 branch of the UPR. Decreased ATF6 activation, as might occur in the aging human population, attenuated the adaptive cardiac hypertrophic growth response to stress and fostered protein misfolding and cardiac dysfunction, highlighting the importance of the UPR for myocardial function in the adult and aging heart during cardiac stress.

### Decrease of the UPR during postnatal development in tissues with low mitotic activity

In many age-related diseases the dysfunction of protein homeostasis leads to the accumulation of misfolded proteins. Moreover, interventions that strengthen the proteostasis network extend the lifespan of invertebrates and mammals and suppresses age-related diseases (Eisenberg et al., 2016; Harrison et al., 2009; Kaushik and Cuervo, 2015; Labbadia and Morimoto, 2014). The proteostasis network in the ER, known as the unfolded protein response (UPR) is activated by ER protein misfolding. Upon activation, the UPR initiates signaling to restore proteostasis and other disrupted cellular conditions, such as metabolic homeostasis or redox status (Han and Kaufman, 2017; Hetz and Papa, 2018). Several studies showed that activation of the UPR promotes longevity in yeast, *C. elegans*, and Drosophila (Burkewitz et al., 2020; Chen et al., 2009; Henis-Korenblit et al., 2010; Kaushik and Cuervo, 2015; Labunskyy et al., 2014; Matai et al., 2019; Sekiya et al., 2017; Shore et al., 2012). Most of these studies were performed in invertebrates and examinations of the proteostasis network during mammalian aging are limited. Recent studies in *C. elegans* have indicated that the collapse of proteostasis is a programmed event that occurs at an early stage of adulthood (Ben-Zvi et al., 2009; David et al., 2010; Labbadia and Morimoto, 2014). To investigate whether a collapse of ER proteostasis occurs as a function of age in mammals, we evaluated the UPR and protein synthesis capacity of neonatal (7 days), young adult (10 weeks), and middle-aged (52 weeks) mice for different mitotic and low-mitotic tissues. Similar to invertebrates, both the UPR and protein synthesis were decreased in young adult mice, especially in tissues with low mitotic activity. Additionally, cardiac gene expression data indicated that the expression of some UPR genes is also reduced in humans as a function of age. One possibility for the reduction of the proteostasis network is that high rates of protein synthesis, which are necessary for rapid cellular differentiation early in life and during cell division, is non-essential as cardiac myocytes enter senescence in the adult heart, leading to diminished proteostasis and a new equilibrium later in life (Eisenberg et al., 2016). Indeed, mitotic tissues that have a high protein synthesis capacity throughout life, such as the liver, retained high levels of the UPR with only a minor reduction throughout adulthood, whereas low-mitotic tissues showed a drastic reduction in protein synthesis and the UPR. A recent study showed that the attenuation of protein synthesis in hematopoietic stem cells increases proteome quality (Hidalgo San Jose et al., 2020), most likely by fostering an adaptive balance between protein synthesis and the UPR. However, whether this occurs in other mammalian tissues remains largely unresolved. Moreover, whether age-related decreases of protein synthesis in adult and aged mammals is a secondary response to an impaired proteostasis, or whether other events trigger the downregulation of protein synthesis and the proteostasis network, remain unknown.

### Responsiveness of the UPR to proteotoxic stress is impaired in adult cardiac myocytes

A reduced proteostasis network in and of itself may not be pathogenic, if it can be increased in response to cellular stress. However, previous studies in *C. elegans* showed that induction of the heat shock response and the UPR is largely compromised after early adulthood (Ben-Zvi et al., 2009; Taylor and Dillin, 2013). While the induction of the UPR was compromised in adult hearts in our study, ATF6 insufficiency did not majorly affect cell survival and UPR expression under baseline conditions in adult hearts, which is coordinate with previous reports (Blackwood et al., 2019a; Correll et al., 2019; Jin et al., 2017). Therefore, ATF6, appears to act primarily as a stress transcription factor in the adult heart, which can be activated by different stimuli, albeit to lower levels than during younger age.

### Loss of ATF6 leads to the accumulation of misfolded proteins in the myocardium and cardiac dysfunction

Recent studies have shed light on the involvement of proteostasis-related signaling pathways in cardiac diseases. However, quite surprisingly, relatively little is known about when, and to what extent protein misfolding occurs in the heart, and whether it significantly contributes to heart disease (Singh and Robbins, 2018). Much of our understanding of the involvement of misfolded proteins in cardiac disease arises from studies of desmin-related (cardio)myopathy, a heterogeneous group of human myopathies which are characterized by the presence of large amounts of intra-sarcoplasmic protein aggregates (Wang and Robbins, 2006). However, even in those cardiomyopathies it remains the subject of ongoing investigation as to whether aggregates are inherently toxic and contribute to myocardial dysfunction or whether they may even be protective (McLendon and Robbins, 2015). Recent findings indeed suggest that the cytotoxic source of misfolded protein are soluble pre-amyloid oligomers, protofibrils, and other intermediates in the amyloid fibril pathway (Singh and Robbins, 2018). Moreover, whether protein misfolding contributes to other cardiac diseases remains to be shown (Singh and Robbins, 2018).

Enhancing protein quality control pathways protects from cardiac disease and protein misfolding (Bhuiyan et al., 2013; Blackwood et al., 2019b; Hofmann et al., 2019; Lynch et al., 2012; Schiattarella et al., 2019; Wang and Robbins, 2006; Yao et al., 2017). Here, we found activation of the ATF6 branch of the UPR in response to acute pressure overload, similar to previous reports (Blackwood et al., 2019a; Lynch et al., 2012). In addition, we showed that protein misfolding can occur in the early phases of heart disease even before cardiac function is compromised. We also showed that UPR is compromised upon early adulthood in the heart. However, in the adult heart, UPR activation was sufficient to protect against protein misfolding and cardiac dysfunction. Nonetheless, it is known that the proteostasis network can collapse during continuous proteotoxic stress, or in aging (Labbadia and Morimoto, 2015). Therefore, it is possible that ongoing proteotoxic stress results in the collapse of the proteostasis network in the adult heart, which may contribute to cardiac disease progression. This would be coordinate with previous studies which found misfolded proteins in (end-stage) heart failure and preclinical studies which showed that enhancing the proteostasis network protects against cardiac diseases. However, this concept remains to be examined in future studies in animal models and humans.

in summary, this study provides insights into the status and functions of the UPR during mammalian postnatal development and the contribution of the ATF6 branch of the UPR to adult myocardial function during cardiac stress. Insufficient UPR activation, due to decreases in ATF6 induced protein misfolding, diminished the hypertrophic cardiac growth response, and impaired myocardial function in response to cardiac stress. Our study suggests that increasing the UPR and, specifically, increasing or activating ATF6 might be a promising strategy to target proteotoxicity-driven heart disease and contributes to the growing body of evidence that protein misfolding directly contributes to cardiac disease progression.

## Methods

### Key Resources Table

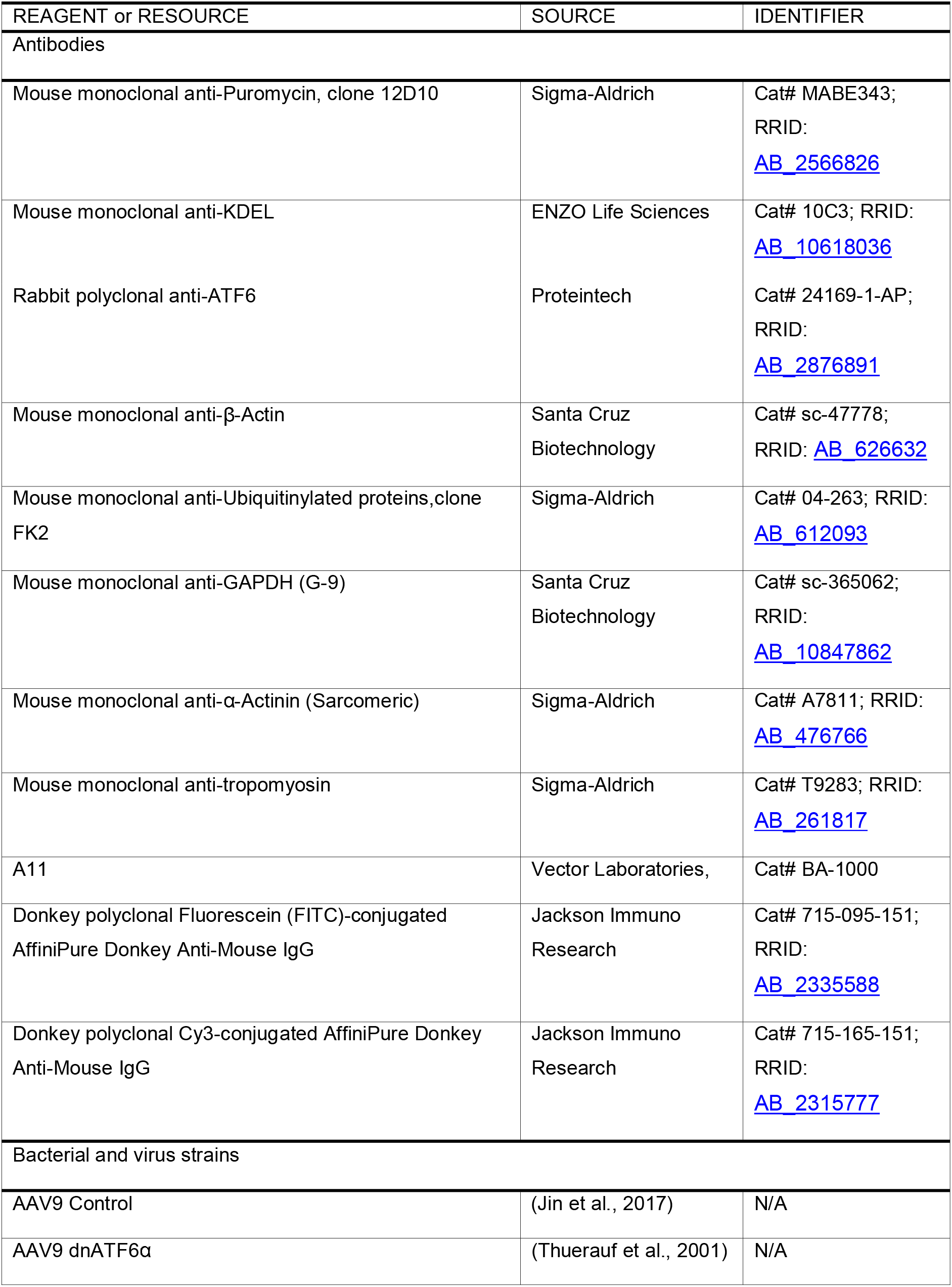
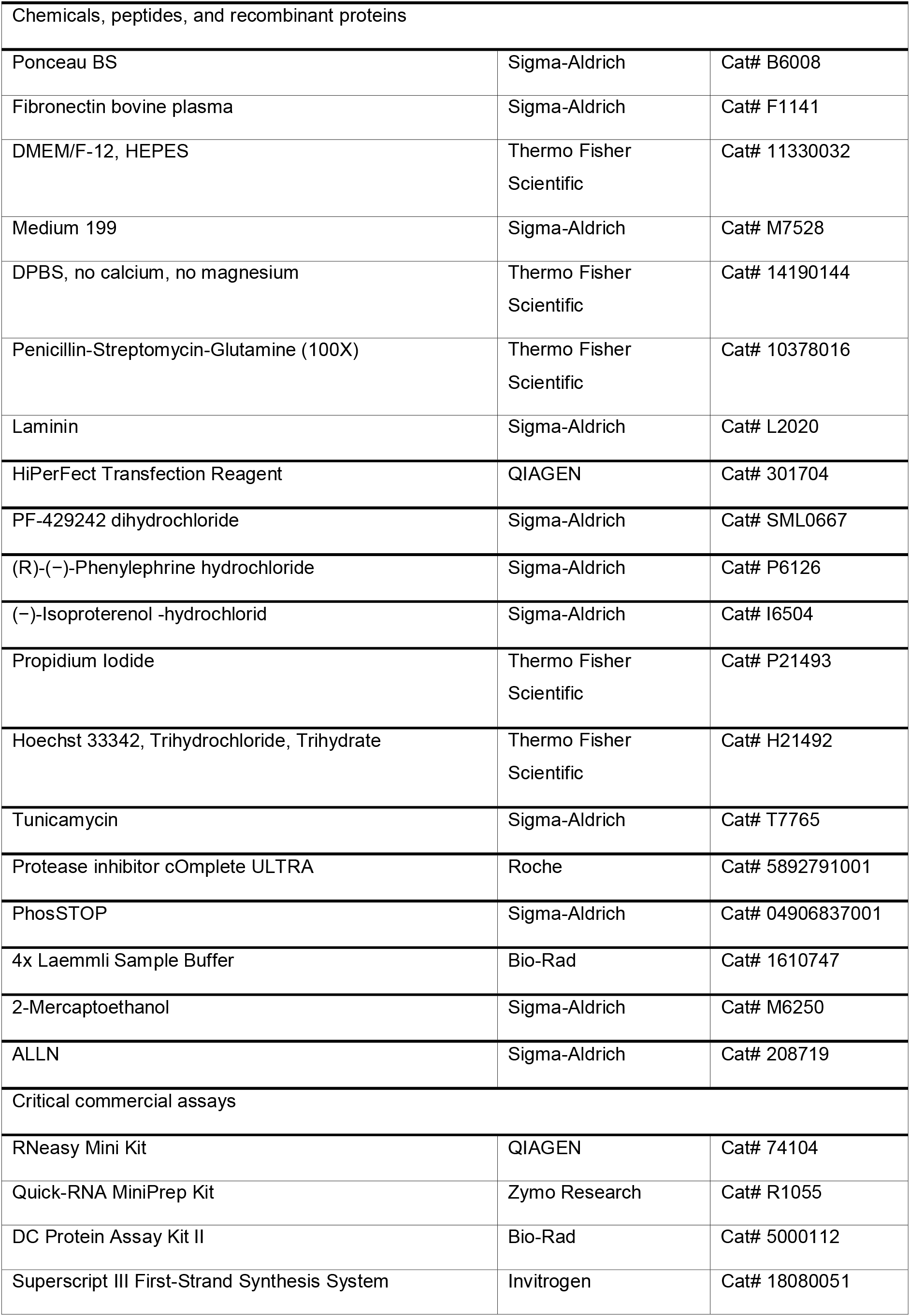
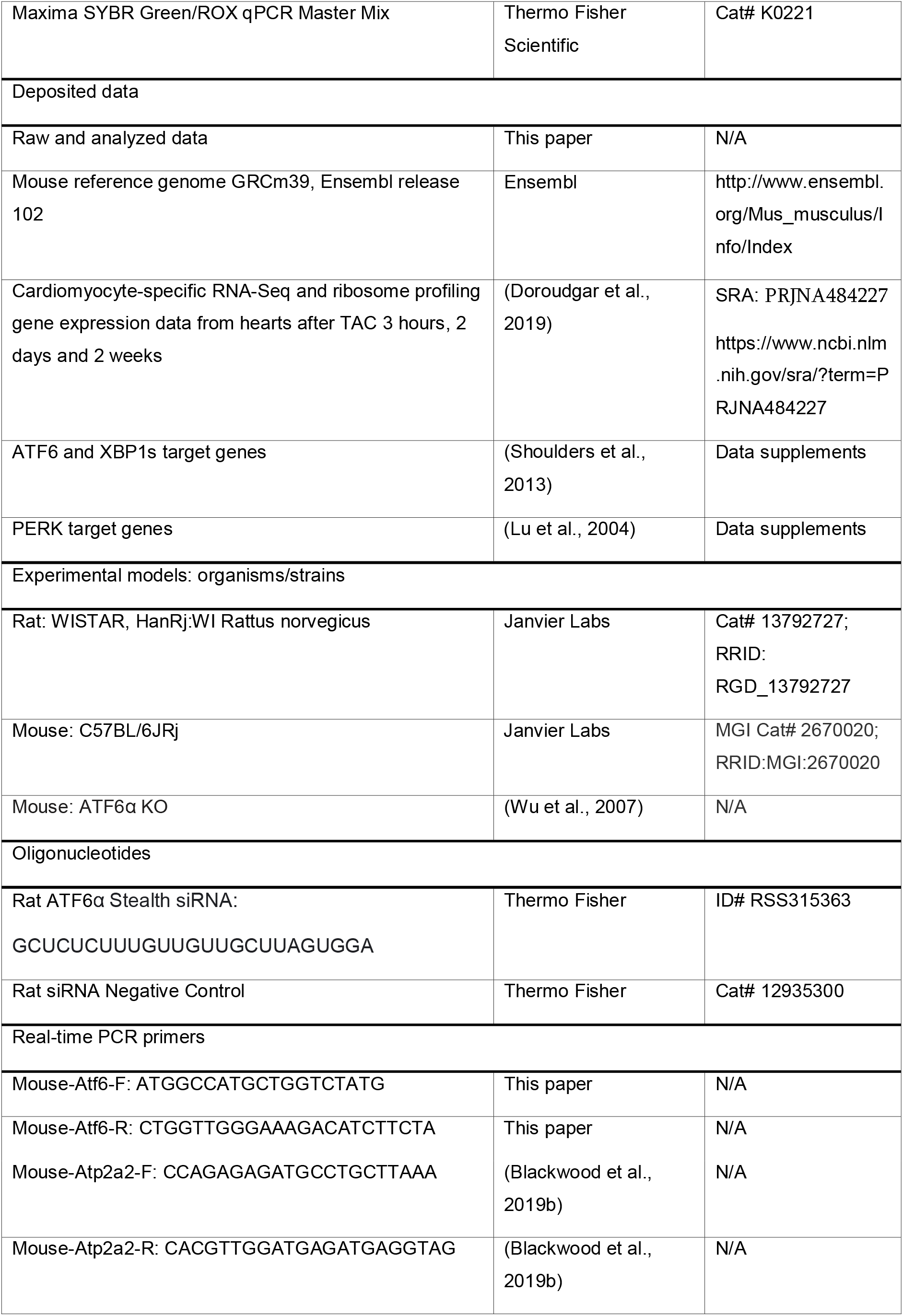
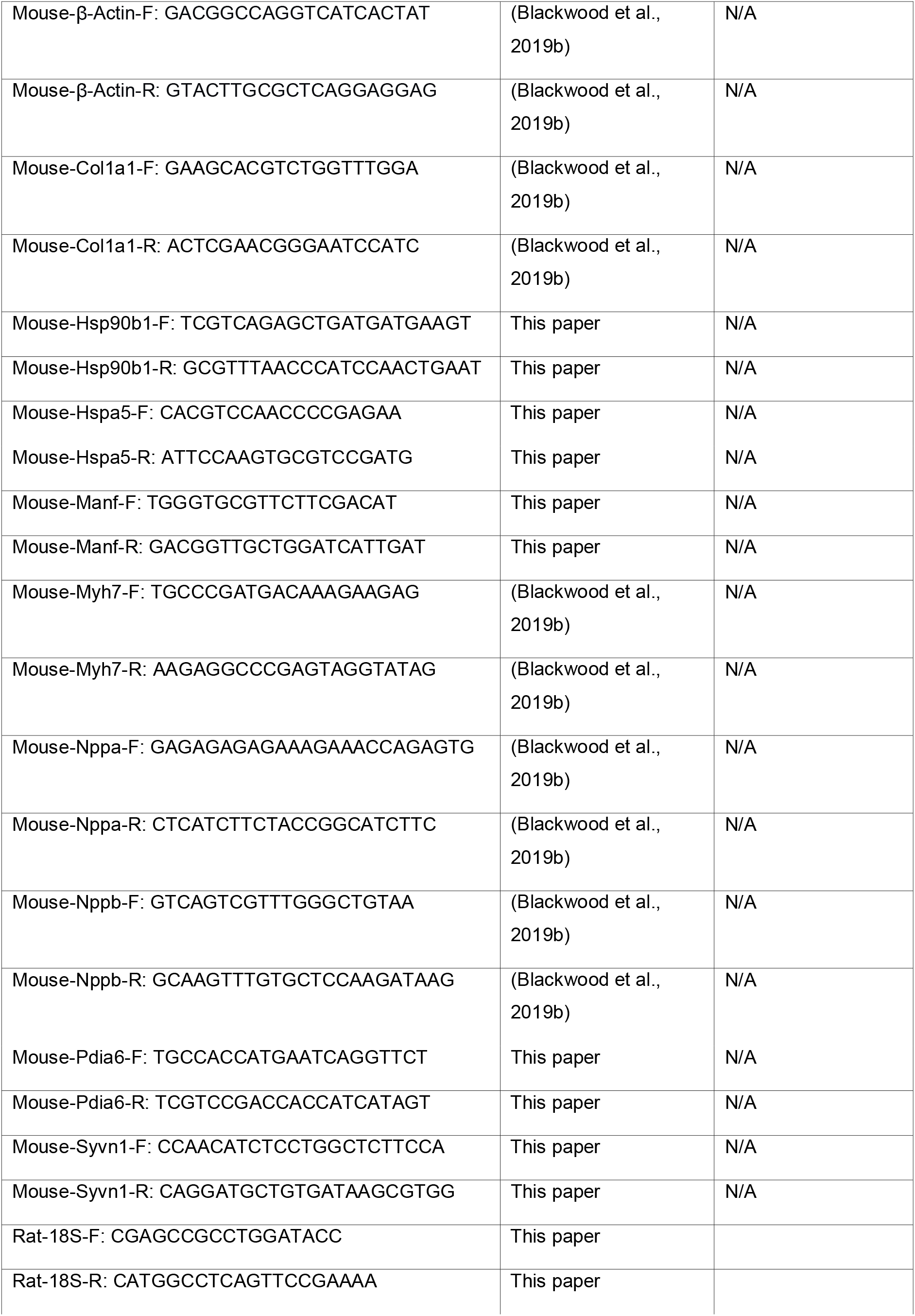
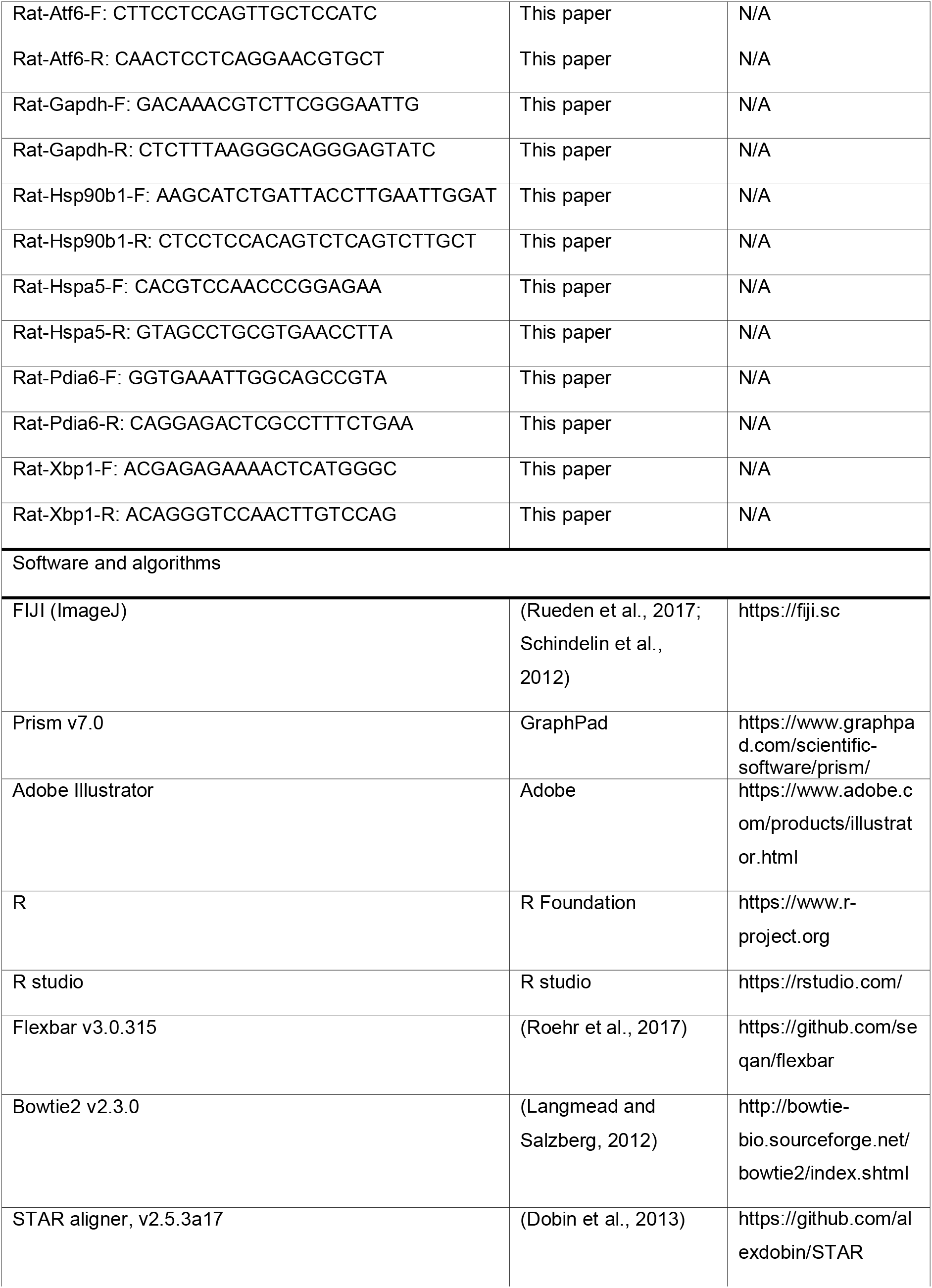
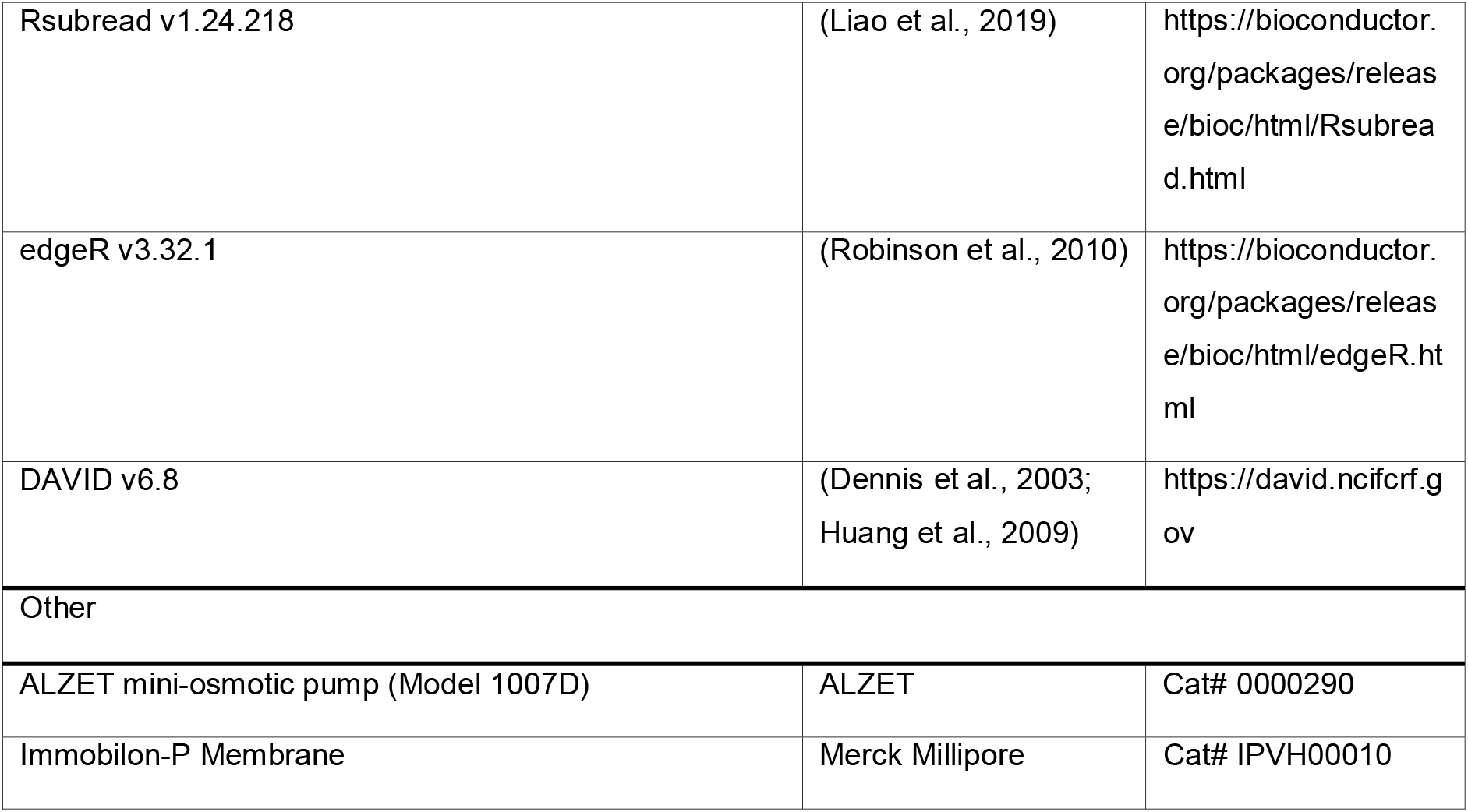

### Resource availability

#### Lead contact

Further information and requests for resources and reagents should be directed to and will be fulfilled by the lead contact, Shirin Doroudgar.

#### Materials availability

This study did not generate new unique reagents.

#### Data and code availability statement

The raw sequencing data generated during this study is available in the NCBI SRA under BioProject ID PRJNA735269. Code used for data analyses is available at https://github.com/jakobilab/Hofmann_et_al_2021.

### Experimental model and subject details

#### Laboratory animals

Data associated with all animal studies reported in this article has been reviewed and approved by the institutional animal care and use committee and conforms to the guide for the care and use of laboratory animals published by the National Research Council. All experiments were performed in 7 day, 10-week and 52-week-old male mice unless otherwise indicated. WT mice used for experiments in which no comparison to ATF6 KO mice was performed were purchased from Jackson Laboratory (Jackson Stock No: 005304). In agreement with previous reports, ATF6 deletion did not affect mouse development (Blackwood et al., 2019a; Jin et al., 2017; Wu et al., 2007). In addition, we did not observe any gender differences on the effects of ATF6 on cardiac structure and function in mice. Due to these observations, and in an attempt to decrease the total number of mice needed to adequately power the study, only male mice were used. All animals were fed *ad libitum* and were housed in a temperature- and humidity-controlled facility with a 12-h light-dark cycle. In the experiments where Puromycin was used to determine protein synthesis rate in vivo, 40mg/kg of puromycin diluted in PBS was given via intraperitoneal injection 30 minutes before animals were sacrificed.

#### Cultured neonatal rat cardiac myocytes (NRVMs)

Isolation of neonatal rat ventricular cardiac myocytes was performed as previously described (Jin et al., 2017). Cells were isolated from one to two-day old Wistar rats (cat# 13792727, WISTAR HanRj:WI, Janvier Labs) via enzymatic digestion and purified by Percoll density gradient centrifugation. Cardiac myocytes were then plated at a density of 0.5 × 10^6^ cells per well on 34.8 mm plastic plates that had been pre-treated with 5 μg/mL fibronectin (cat# F1141, Sigma-Aldrich) in serum free DMEM/F12 medium (cat# 11330032, Thermo Fisher Scientific) for one hour and then cultured in DMEM/F12 1:1, containing 10% fetal bovine serum, 100 units/mL of penicillin, 100μg/mL streptomycin and 292 μg/mL glutamine (cat# 10378016, Thermo Fisher Scientific). After 24h, media was changed to DMEM/F12 1:1 supplemented with BSA (1 mg/mL), 100 units/mL of penicillin, 100μg/mL streptomycin and 292 μg/mL glutamine or when neonatal cardiac myocytes were directly compared to adult cardiac myocytes to DMEM/F12 1:1, containing only 100 units/mL of penicillin, 100μg/mL streptomycin and 292 μg/mL glutamine for 24h after which the cells were subjected to the respective treatments.

#### Cultured adult rat ventricular cardiac myocytes (ARVMs)

Adult rat ventricular myocytes were isolated from 6-week old male Wistar rats (WISTAR RjHan:WI, Janvier Labs) via enzymatic digestion as previously described (Jin et al., 2017; Völkers et al., 2011). Hearts were rapidly excised and perfused with a rate of 8 mL/min in a Langendorff apparatus. After excision, hearts were initially perfused with calcium-free medium (ACM) (pH 7.2) consisting of (in mM) 5.4 KCl, 3.5 MgSO4, 0.05 pyruvate, 20 NaHCO3, 11 glucose, 20 HEPES, 23.5 glutamate, 4.87 acetate, 10 EDTA, 0.5 phenol red, 15 butanedionemonoxime (BDM), 20 creatinine, 15 creatine phosphate (CrP), 15 taurine and 27 units/mL insulin under continuous equilibrium with 95% O2 and 5% CO2. After 5 min the perfusion was switched to ACM plus collagenase (0.5 U/mL) for 20 – 30 min. Finally, perfusion was changed to low Na+, high sucrose Tyrode solution containing (in mM) 52.5 NaCl, 4.8 KCl, 1.19 KH2PO4, 1.2 MgSO4, 11.1 glucose, 145 sucrose, 10 taurine, 10 HEPES, 0.2 CaCl2 for 15 min. Thereafter left ventricles of digested hearts were cut into small pieces and subjected to gentle agitation to allow for dissociation of cells. Consequently, cells were resuspended in ACM without BDM in which 2 mM extracellular calcium was gradually reintroduced at 25 °C. ARVMs were cultured on laminin coated culture dishes. Cells were plated in Medium 199 (cat# M7528, Sigma-Aldrich) supplemented with 100 units/mL of penicillin, 100μg/mL streptomycin and 10μg/mL laminin (cat# L2020, Sigma-Aldrich). After one hour, media was changed to M199 media supplemented with 100 units/mL penicillin, 100μg/mL streptomycin. Cells were subjected to the respective treatments after 24h. To induce ER stress *in vitro*, 10 μg/mL tunicamycin (TM) was added to the cell media for 24h in the indicated experiments. Other concentrations that were used are described in the respective figures.

### Method details

#### siRNA transfection

Neonatal rat ventricular myocytes were plated at a density of 0.5 × 10^6^ cells per well on fibronectin coated 34.8 mm plates and cultured in Dulbecco’s modified Eagle’s medium (DMEM)/F12 1:1 (cat# 11330-032, Thermo Fisher Scientific), containing 10% fetal bovine serum, 100 units/mL of penicillin, 100μg/mL streptomycin and 292 μg/mL glutamine for 24h. One day after, media was changed to DMEM/F12 1:1 containing 0.5% fetal bovine serum, 120 nM of siRNA and 0.625% HiPerFect (cat# 301704, QIAGEN) and no antibiotics. HiPerFect transfection reagent was used to transfect cells with small interfering (si) RNA oligoribonucleotides targeted to rat ATF6 (assay ID# RSS315363, Stealth siRNA, Thermo Fisher). A non-targeting sequence (cat# 12935300, Thermo Fisher) was used as a control siRNA. NRVCM were incubated with the siRNA for 16 hours. Then, medium was changed to DMEM/F12 1:1 supplemented with BSA (1 mg/mL), 100 units/mL of penicillin, 100μg/mL streptomycin and 292 μg/mL glutamine. Respective treatments were initiated after 48 hours.

#### Adenovirus generation and transfection

Plasmid vector construction and recombinant adenovirus generation encoding AAV9-control, AAV9-3XFLAG-tagged constitutively active ATF6, ATF6α(1-373), and dominant-negative ATF6α (dnATF6) has been described previously (Jin et al., 2017; Thuerauf et al., 2001; Wang et al., 2000). Transduction was performed by incubating cultures overnight with the appropriate adenovirus at a multiplicity of infection (MOI, number infectious particles per cell) of five. On the next day media was changed to DMEM/F12 1:1 supplemented with BSA (1 mg/mL), 100 units/mL of penicillin, 100μg/mL streptomycin and 292 μg/mL glutamine. After 24h, desired treatments were initiated.

#### Cultured cell area

Phase-contrast microscopy pictures were taken from five different fields per culture (n=3 individual cell cultures for each treatment). Cell area for at least 500 cells per condition was determined using FIJI (Rueden et al., 2017; Schindelin et al., 2012). The number of measured cells can be found in the respective figure legends. For cultured myocytes cell area, myocytes were seeded at a density of 0.5 × 10^6^ cells per well on fibronectin coated 34.8 mm plates maintained in Dulbecco’s modified Eagle’s medium (DMEM)/F12 1:1, containing 10% fetal bovine serum, 100 units/mL of penicillin, 100μg/mL streptomycin and 292 μg/mL glutamine for 24h. After 24h media was changed to DMEM/F12 1:1 supplemented with BSA (1 mg/mL), 100 units/mL of penicillin, 100μg/mL streptomycin and 292 μg/ml glutamine. After an additional 24 hours, cells were pretreated with either DMSO or 50μM of PF-429242 dihydrochloride (cat# SML0667, Sigma-Aldrich), diluted in DMSO, for 30 minutes and then treated with Dulbecco’s phosphate buffered saline (PBS) (cat# 14190-144, Thermo Fisher Scientific), (R)-(-)-phenylephrine hydrochloride (50μM) (cat# P6126, Sigma-Aldrich) or isoproterenol (10μM) diluted in PBS, for 24h. For siRNA and adenovirus transfection cells were treated as described above.

#### Immunofluorescence

Cardiac myocytes were isolated as described above. Cardiac myocytes were then plated on glass chamber slides that had been pre-treated with 25 μg/mL fibronectin for one hour. After the respective treatments, the slides were washed twice in PBS and then fixed with 4% PFA for 20 minutes. Slides were washed three times for 5-10 minutes in PBS and then incubated for 10 minutes in permeabilization buffer consisting of PBS, 0.2% Triton X-100 and 0.1M glycine. Then slides were washed once with PBS and then incubated with blocking buffer containing 10% horse serum and 0.2% Triton X-100 in PBS for one hour. Next, slides were incubated overnight at 4°C with primary antibody diluted in blocking buffer. The primary antibody used was anti-α-actinin (sarcomeric) (cat# A7811, Sigma-Aldrich; 1:100). On the next day slides were washed three times for 5-10 minutes in PBS and then incubated with secondary antibody in blocking buffer for one hour. The secondary antibody used was fluorescein (FITC)-conjugated AffiniPure donkey anti-mouse IgG (cat# 715-095-151, Jackson Immuno Research; 1:100). The slides were then washed three times for 5-10 minutes in PBS and incubated in a 1:500 solution of 1mM TO-PRO-3 iodide (cat# T3605, Thermo Fisher Scientific) diluted in PBS for 10 minutes. Then slides were washed twice in PBS for 10 minutes and covered with VECTASHIELD Antifade Mounting Medium (cat# H-1000, Vector Laboratories) and a glass plate and visualized by microscopy. Immunocytofluorescence of mouse heart sections was performed as previously reported (Doroudgar et al., 2015). Primary antibodies used were anti-tropomyosin (cat# T9283, Sigma-Aldrich; 1:200) and anti-A11 (1:200). For detection of A11, a biotinylated goat anti-rabbit (Vector Laboratories, BA-1000) secondary was required. Secondary antibodies used were Alexa Fluo 488. Images were obtained using a Leica DMi8 confocal laser scanning microscope.

#### Cell viability

For quantification of viability cells were subjected to the respective treatments. Then, 1 μg/mL of propidium iodide (cat# P21493, Thermo Fisher Scientific) and 5 μg/mL of Hoechst 33342, trihydrochloride, trihydrate (cat# H21492, Thermo Fisher Scientific) was added to the culture media. After 10 minutes fluorescent images of random positions of the cell culture area were taken using a ZEISS Axio Vert.A1 microscope. Propidium iodide positive and negative cells were quantified with FIJI (Schindelin et al., 2012).

#### Isoprenaline model of cardiac hypertrophy

Male WT and ATF6 KO mice at 10 weeks of age were randomly assigned to the experimental groups. To induce cardiac hypertrophy, mice were subjected to either Lactated Ringer’s solution or 30mg/kg/day Isoprenaline (cat# I6504, Sigma-Aldrich) treatment via ALZET mini-osmotic pump implants (Model 1007D, cat# 0000290) for seven days. For osmotic pump implantation, mice were anesthetized with isoflurane. Pumps were inserted subcutaneously through a small incision at the lower back. Mice received daily intraperitoneal injections of 1μl/g of 0.1 mg/mL buprenorphine solution, diluted in phosphate-buffered saline at time of surgery and up to 2d after surgery.

#### TAC surgery

Animals were randomly assigned to the experimental groups. TAC was performed as previously described (Blackwood et al., 2019a). Briefly, adult male mice were anesthetized using a 2% isofluorane/O2 mixture and intubated. Mice were treated with buprenorphine (0.1 mg/kg IP) and a partial trans-sternal thoracotomy performed using aseptic technique. An approximately 1.5 cm vertical left parasternal skin incision was made, underlying pectoralis muscle retracted, and the chest cavity entered through the fourth intercostal space. Using hooked retractors, adjacent ribs were retracted to expose the heart and aortic arch. The aorta was isolated from annexed tissue, and the artery partially ligated between the innominate and left common carotid arteries with 6-0 silk. The calibrated constriction of the aorta was performed by placing a dull 27-gauge needle to the side of the artery, the ligature tied firmly to both the needle and the artery, and, subsequently, the needle was removed leaving a calibrated stenosis of the aorta. Sham operated mice were exposed to the same procedure, except that the aorta was not constricted. The thoracic cavity was closed and the animals were allowed to recover. Animals were injected once with buprenorphine (0.1 mg/kg IP) about 12 h after recovery in order to reduce any post-operative discomfort. In case any animals displayed signs of pain or distress after this period, additional doses of buprenorphine were administered as needed. Sham-operated mice were euthanized at time points matching to TAC surgery time points.

#### Echocardiography

Echocardiography was carried out on anesthetized mice using a Visualsonics Vevo 2100 high-resolution echocardiograph (Tsujita et al., 2005). Anesthesia was administered via a facial mask and maintained by a minimum dose of isoflurane (1.0–2.0%). Echocardiography was performed at a heart rate of 450-550 bpm.

#### Preparation of tissue lysates

Mice were sacrificed and different tissues were rapidly excised, washed in PBS and snap frozen in liquid nitrogen. Different tissues were homogenized using a tissue homogenizer in 5 volumes of ice-cold polysome buffer containing 20mM Tris pH 7.4, 10mM MgCl, 200mM KCl and 1% Trition X-100. For protein analysis via immunoblotting, initial lysates were further diluted with 9 volumes of RIPA buffer containing 20mM Tris-HCl (pH 7.4), 150mM NaCl, 1% Triton X-100, 0.1% SDS, 0,5% Sodium deoxycholate, protease inhibitor cOmplete ULTRA (cat# 05892791001, Roche) and phosphatase inhibitor PhosSTOP (cat# 04906837001, Sigma-Aldrich). RNA was isolated from tissue lysates using the RNeasy Mini Kit (cat# 74104, Qiagen).

#### Immunoblotting

Cultured cells and mouse tissues were lysed in RIPA Buffer consisting of 50mM Tris pH 7.5, 150mM NaCl, 1% Triton X-100 and 1% SDS, which was supplemented with protease inhibitor cOmplete ULTRA (cat# 05892791001, Roche) phosphatase inhibitor PhosSTOP (cat# 04906837001, Roche). Lysates were cleared by centrifugation at 4 °C for 10 minutes at 20.000 rcf. Lysate protein concentration was determined using DC Protein Assay Kit II (cat# 5000112, Bio-Rad). Equivalent amounts of protein, usually 20-30 μg, were brought up to similar volume, mixed with Laemmli Sample Buffer (Bio-Rad; 161-0747) and 2-Mercapoethanol (cat# M6250, Sigma-Aldrich) and boiled at 95°C for 5 minutes. Samples were separated on SDS-PAGE gels and transferred to Immobilon-P transfer membranes (cat# IPVH00010, Merck Millipore). The following antibodies were used to probe the membranes: GAPDH (cat# G-9, sc-365062, Santa Cruz Biotechnology; 1:20,000), β-Actin (cat# C4, sc-47778, Santa Cruz Biotechnology; 1:20,000), KDEL (cat# 10C3, ADI-SPA-827, ENZO Life Sciences; 1:2,000-5,000), ATF6 (cat# 24169-1-AP, Proteintech; 1:1,000), Puromycin (cat# MABE343, Merck Millipore; 1:2,000-1:10,000), anti-Ubiquitinylated proteins,clone FK2 (cat# 04-263, Merck Millipore; 1:1000). Ponceau solution was prepared with Ponceau BS (cat# B6008, Sigma Aldrich).

#### Quantitative Real Time PCR

Total RNA was isolated from cultured cardiomyocytes using the Quick-RNA MiniPrep Kit (cat# R1055, Zymo Research) and from tissue using the RNeasy Mini Kit (cat# 74104, Qiagen). cDNA was generated by reverse transcription using Superscript III First-Strand Synthesis System (Invitrogen; 18080-051). Quantitative Real Time PCR was performed with Maxima SYBR Green/ROX qPCR Master Mix (Thermo Fisher cat# K0222) in a StepOnePlus RT-PCR System (Thermo Fisher).

#### Xbp1 splicing

XBP1 splicing was assessed with Real Time PCR using the following primers: FW ACGAGAGAAAACTCATGGGC; Rev ACAGGGTCCAACTTGTCCAG. cDNA was run on a 2% agarose gel at 130V for one hour and viewed by UV illumination.

#### Sequencing data analysis

Transcript profiling of ATF6 KO mice was performed as previously described. Briefly, total RNA was isolated from mouse left ventricular extracts and RNA sequencing was carried out on Illumina Nextseq at CellNetworks Deep Sequencing Core Facility at Heidelberg University. Sequencing adapter residues and low-quality bases were removed from raw sequencing reads prior to all other analysis steps using Flexbar version 3.0.315 (Roehr et al., 2017). Subsequently, reads mapping to known ribosomal RNA species were excluded from further analyses using Bowtie2 version 2.3.0 (Langmead and Salzberg, 2012) with a custom rRNA index and only keeping non-aligning reads. Principal read mapping against the mouse reference genome (GRCm39, Ensembl release 102) was performed with the STAR aligner, version 2.5.3a17 (Dobin et al., 2013). The read-to-transcript assignment was carried out using the R package Rsubread version 1.24.218 (Liao et al., 2019) and the ENSMBL gene annotation GRCm39, Ensembl release 102. The resulting count table was further processed with the edgeR R package19 (Robinson et al., 2010) to construct the list of differentially expressed genes. RNA-Seq, ribosome profiling and proteomics gene expression data of TAC 3h, 2d, and 2wk relative to time-matched Sham mice was obtained from our previously published dataset (Doroudgar et al., 2019). Significantly regulated genes were defined as FDR <0.05. ATF6 and XBP1s target genes were identified from Shoulders et al. (2013). Due to incorrectly reported q-values in the available gene list from Shoulders et al. (2013) (Shoulders et al., 2013), novel q-values were derived from the reversed log10 scores of Benjamini-Hochberg. PERK-selective target genes were identified from Lu et al. (Lu et al., 2004). PERK-selective target genes were defined as genes induced both >2 fold in AP20187 treated eIF2a (S/S) Fv2E-PERK cells at 8h and >2 fold in TM treated WT cells at 6 hours. ‘Unfolded protein response genes’ were defined as the merged gene entries from the GO categories GO:0006986 (response to unfolded protein), GO:0030968 (endoplasmic reticulum unfolded protein response) and GO:0034976 (response to endoplasmic reticulum stress).

#### Gene Ontology analysis

GO term enrichment analysis was performed as previously described. Briefly, genes with FDR <0.05 were considered for further analysis. GOTERM_BP_DIRECT in DAVID v6.8 (Dennis et al., 2003; Huang et al., 2009) was used with the subset of expressed protein-coding genes as background set. Only enriched GO terms with at least five significantly changed genes were kept for further analysis and ranked by p-value. Top enriched terms were retained and visualized with a custom plotting routine showing enrichment p-value.

#### Quantification and statistical analysis

Statistical analysis was performed using GraphPad Prism 7.0 (Graphpad Software Inc; www.graphpad.com) or R (R Foundation; https://www.r-project.org). Data values are mean ± standard error of the mean (SEM). For statistical analysis one-way ANOVA with Turkey post-hoc analysis or when only two conditions where compared, unpaired two tailed t-test was used. p < 0.05 was defined as significant difference. Details of the statistical analyses of the sequencing data can be found in the method details. Biological replicate numbers for each figure can be found in the accompanying figure legend.

## Acknowledgments

This work was supported by grants from the German Centre for Cardiovascular Research (DZHK), the University of Arizona College of Medicine – Phoenix, and the Translational Cardiovascular Research Center at the University of Arizona College of Medicine – Phoenix to S.D. C.H. acknowledges support by a Boehringer Ingelheim Fonds Travel Grant, the DZHK mobility program, and the German Heart Foundation (Deutsche Herzstiftung). R.J.K. acknowledges support from National Institutes of Health (NIH) grant R01 AG062190. C.C.G. acknowledges NIH grants 1HL135893, 1HL141463, and 1HL149931. E.A.B acknowledges NIH grant 1F31HL140850, the Inamori Foundation, the Achievement Rewards for College Scientists Foundation, Inc, San Diego Chapter, the Rees-Stealy Research Foundation Phillips Gausewitz, MD, Scholarship of the San Diego State University Heart Institute.

## Author contributions

Conceptualization, C.C.G., C.H. and S.D.; Methodology, C.H., C.S., E.A.B., J.G., M.V., N.H., S.D. and T.J.; Investigation C.H., C.S., E.A.B., J.G. and N.H.; Formal Analysis C.H. and T.J.; Visualization C.H. and S.D.; Providing critical reagents and advice R.J.K; Writing – Original Draft C.H. and S.D.; Writing – Review & Editing C.H. and S.D. with input from all authors; Resources C.C.G., H.A.K, M.V. and S.D.; Supervision C.C.G., M.V., S.D.; Funding Acquisition C.C.G., H.A.K., M.V. and S.D.; Project Administration S.D.

