## Supplementary Material for "Age-related decline of the unfolded protein response in the heart promotes protein misfolding and cardiac pathology"

**Table of Contents**

**Supplementary Figure Legends**	**2**

**Supplementary Figures**	**3**

**Supplementary Tables**	**8**

**Supplementary Figure Legends**

**Figure S1. Reduction of ATF6 activity in adult and aged cardiac myocytes.**

(A) Puromycin immunoblot and respective quantification of neonatal and adult cardiac myocytes under baseline conditions. (B) p90 and p50 ATF6 immunoblots and respective quantification of neonatal and adult cardiac myocytes under baseline conditions. 25ug/mL ALLN was added 3h before the end of the experiment. (C) Immunoblots and respective quantifications for GRP94, GRP78, and PDIA6 in neonatal and adult cardiac myocytes under baseline conditions. * indicates p<0.05 from control.

**Figure S2. Induction of the UPR in adult cardiac myocytes in response to ER stress.**

(A and B) *Hspa5* and *Hsp90b1* levels in adult cardiac myocytes treated with indicated doses of TM for 24h. (C) Immunoblot and respective quantifications of GRP94, GRP78, and PDIA6 of adult cardiac myocytes treated with indicated doses of TM for 24h. * indicates p<0.05 from control.

**Figure S3. XBP1 splicing in response to acute ATF6 inhibition.**

(A) RT-PCR using primers that recognize the unspliced and spliced isoform of XBP1. Neonatal cardiac myocytes were treated with indicated doses of the ATF6 inhibitor PF-429242 for 24h. The respective quantification is shown on the right. * indicates p<0.05 from control.

**Figure S4. Transcriptional UPR induction in response to transverse aortic constriction.**

(A, B and C) Volcano plots of transcriptional changes after 3h, 2d or 2wk TAC surgery in mice compared to time-matched sham controls. The highlighted colors indicate ATF6 (red), XBP1s (blue), or PERK (yellow) target genes. The horizontal dotted line indicates an FDR of 0.05. Genes above the line are considered significant (FDR < 0.05). (D) Enrichment of gene ontology terms (biological process) of significantly changed ATF6 (red), XBP1s (blue), or PERK (yellow) target genes 2d after TAC surgery. (E) Heat map showing the transcriptional regulation of all ATF6 (red), XBP1s (blue), or PERK (yellow) target genes 2d after TAC surgery.

**Figure S5. Translational UPR induction in response to transverse aortic constriction.**

(A, B and C) Volcano plots of translational changes after 3h, 2d, or 2wk TAC surgery in mice compared to time-matched sham controls. The highlighted colors indicate ATF6 (red), XBP1s (blue), or PERK (yellow) target genes. The horizontal dotted line indicates an FDR of 0.05. Genes above the line are considered significant (FDR < 0.05). (D) Enrichment of gene ontology terms (biological process) of significantly changed ATF6 (red), XBP1s (blue), or PERK (yellow) target genes 2d after TAC surgery. (E) Heat map showing the translational regulation of all ATF6 (red), XBP1s, (blue) or PERK (yellow) target genes 2d after TAC surgery.

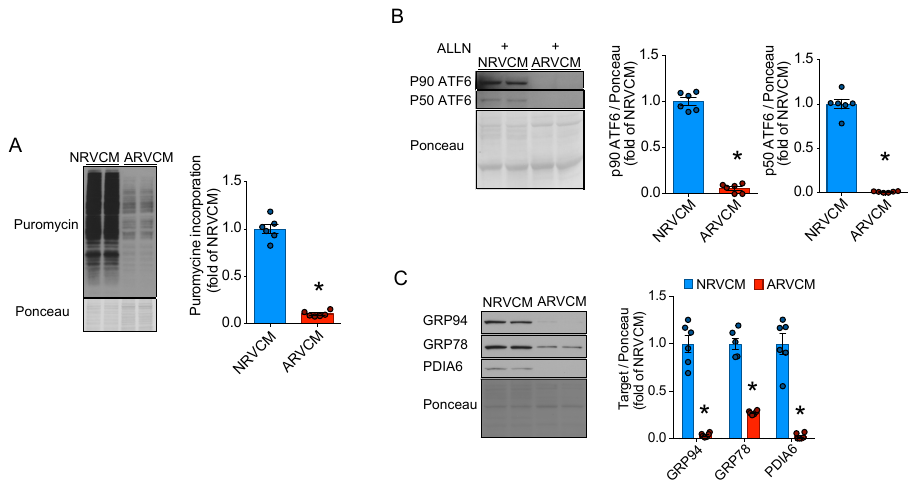

**Supplementary Figure 1**

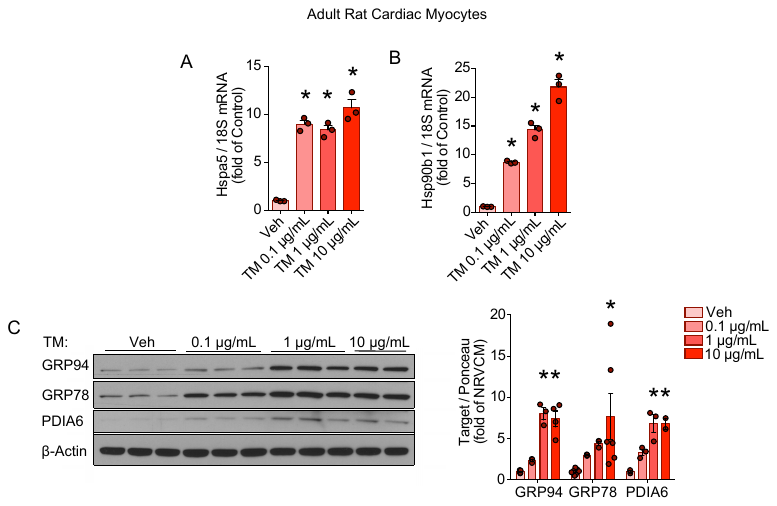

**Supplementary Figure 2**

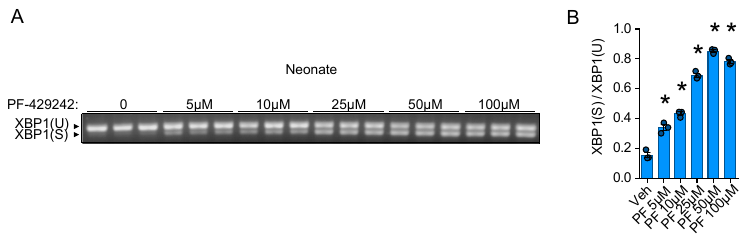

**Supplementary Figure 3**

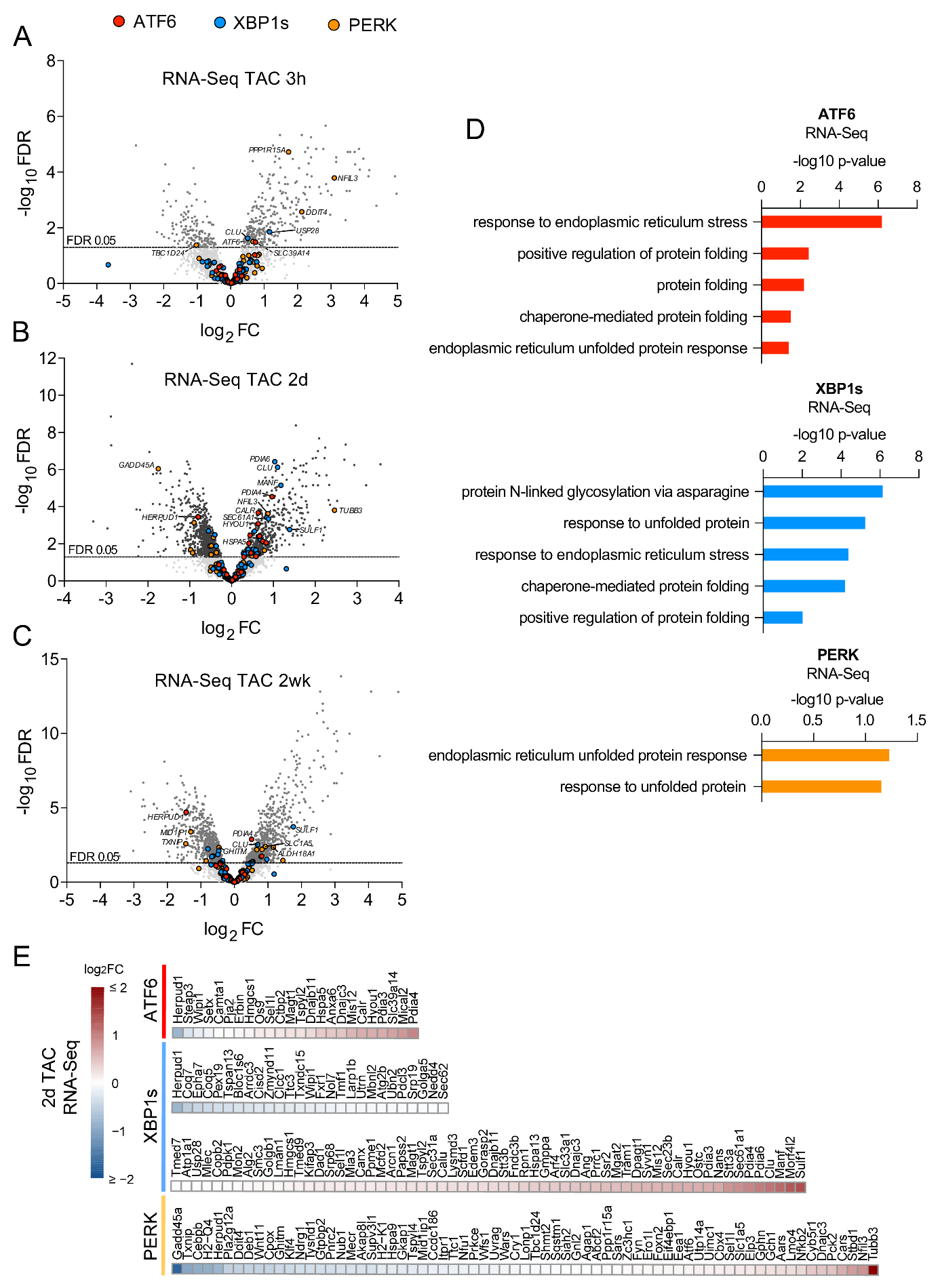

**Supplementary Figure 4**

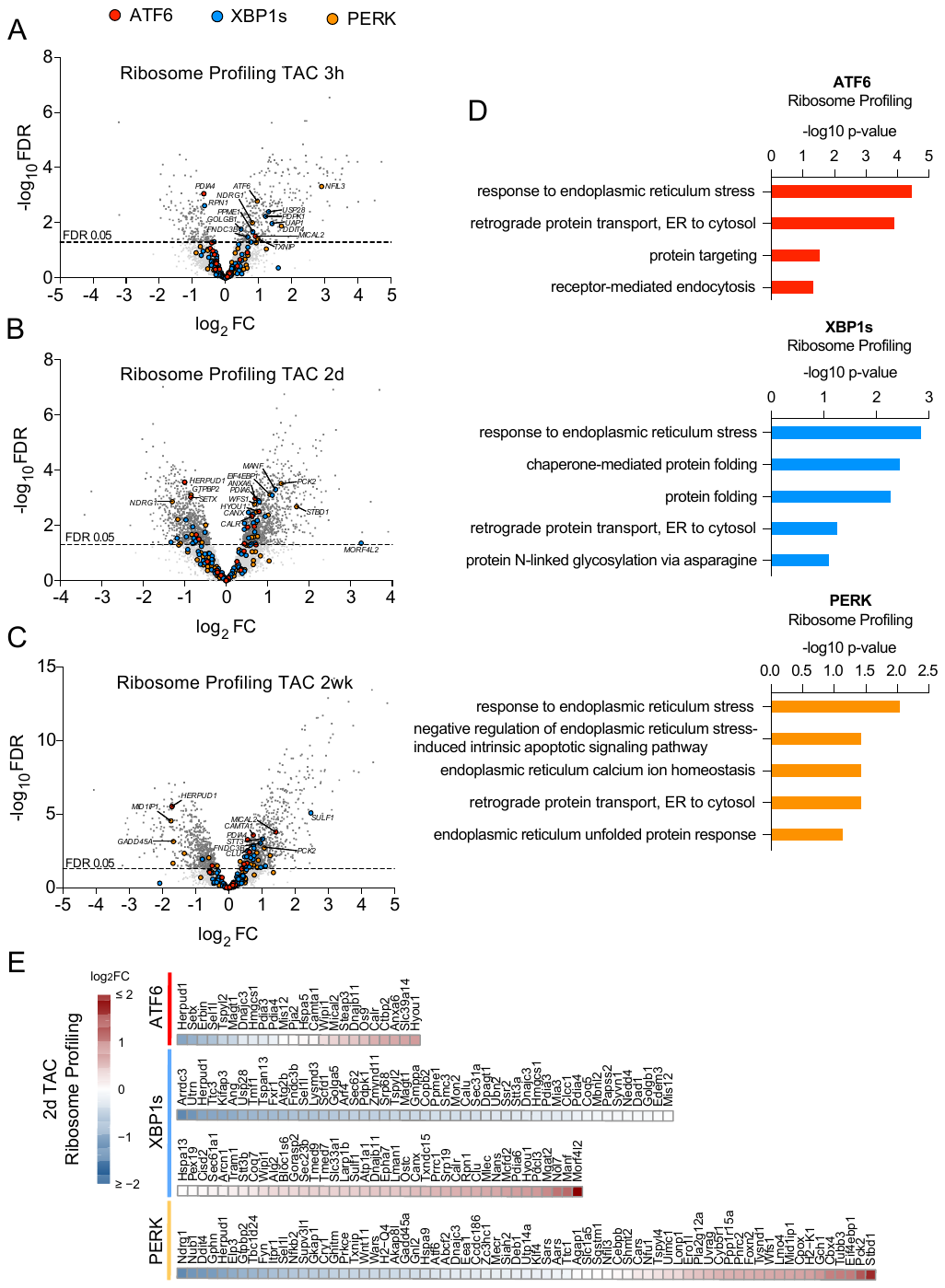

**Supplementary Figure 5**

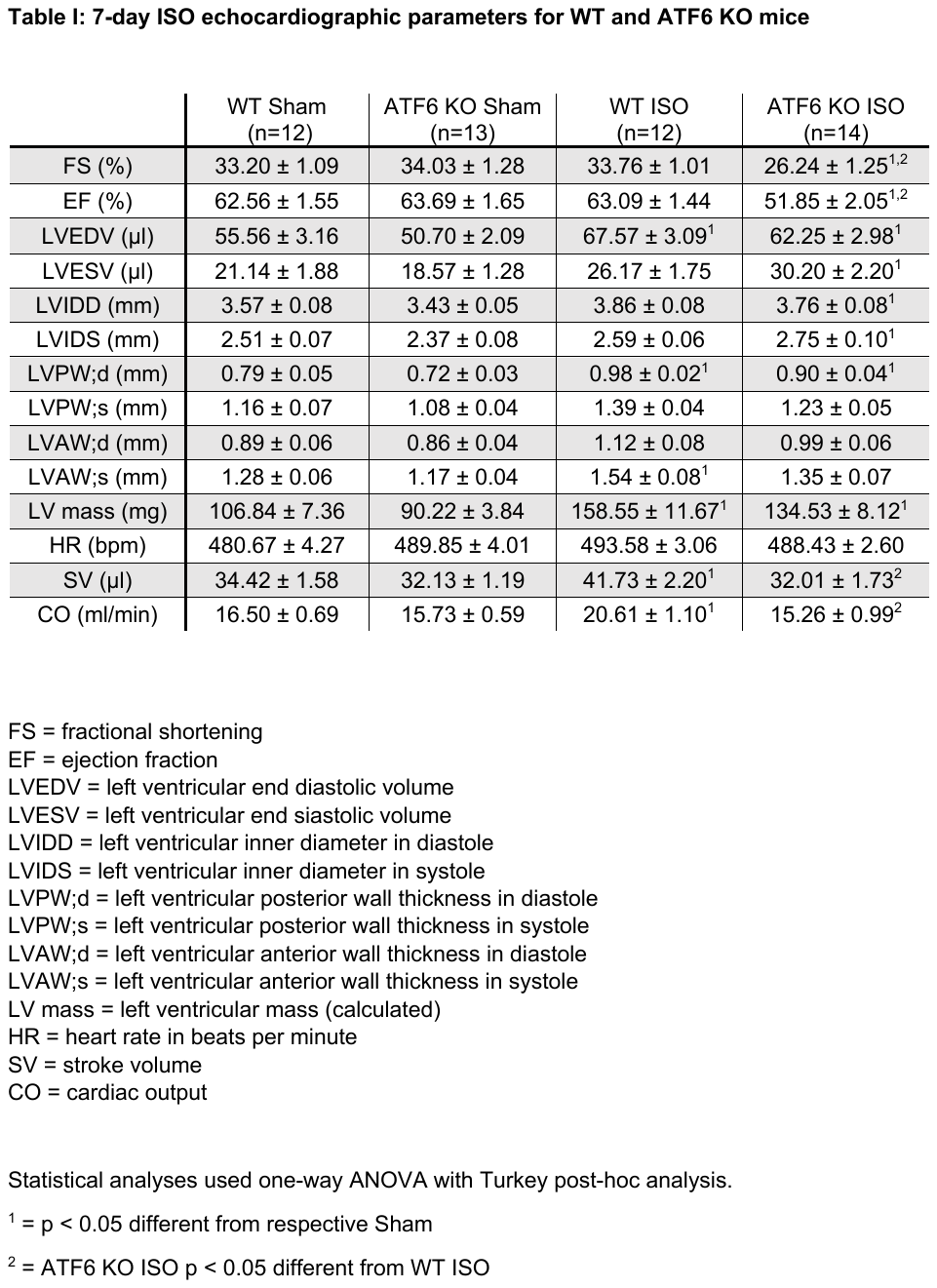

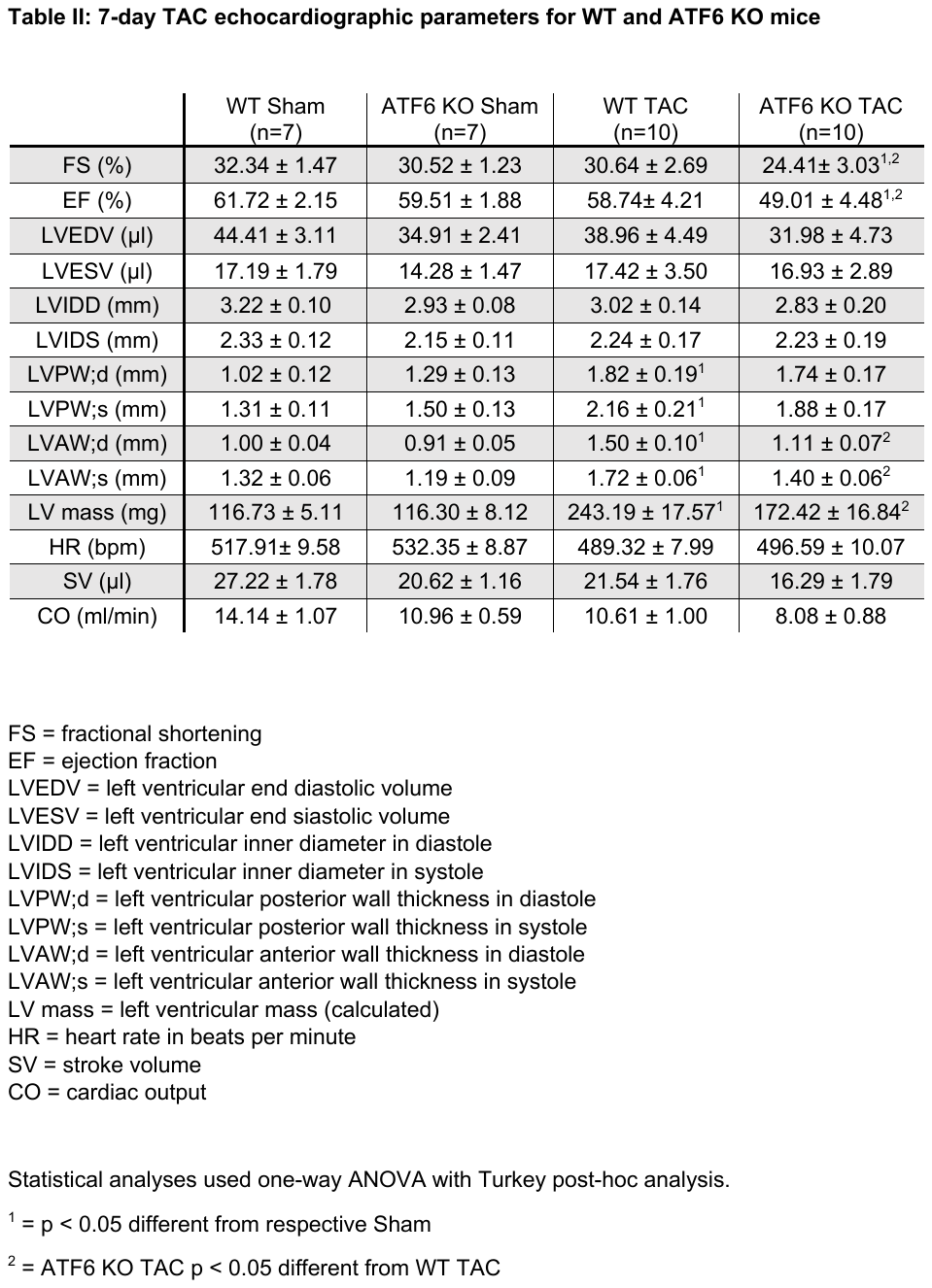
